# Multi-resolution Spatial Graphical Regression Models for Hierarchical Spatial Transcriptomics Data

**DOI:** 10.64898/2026.05.12.724724

**Authors:** Liying Chen, Satwik Acharyya, Allison M. May, Aaron M. Udager, Evan T. Keller, Veerabhadran Baladandayuthapani

## Abstract

Advances in spatial transcriptomics (ST) technologies enable systematic molecular characterization of tumor microenvironment, tumor gradients and gene regulatory networks. Cancer progression is known to vary along pathological gradients, yet existing network approaches for gene network inference typically ignore hierarchical spatial organization across the tumor. We develop a Bayesian multi-resolution spatial graphical regression (`mSGR`) framework to infer spatially varying gene networks from multi-resolution ST data. The proposed model allows precision matrices to vary across hierarchically structured spatial domains, capturing both local and global organization within the tumor. To identify spatially varying regulatory relationships, we introduce a spatially structured edge selection strategy that borrows strength across regions according to spatial proximity and pathological gradients, while Gaussian-process priors flexibly model spatial variation in edge strengths. Scalable inference is achieved through an augmented mean-field variational Bayes algorithm with node-wise parallel regressions, enabling efficient estimation in high-dimensional settings. Simulation studies demonstrate improved recovery of network structures compared with competing approaches. Applying `mSGR` to multi-resolution ST data from kidney cancer reveals stronger regulatory connectivity in transitional regions of epithelial-mesenchymal transition pathway and identifies hub genes along the tumor gradient, illustrating how spatially resolved network analysis can provide key insights into tumor microenvironment organization.

## 1 Introduction

### Scientific background and rationale

The tumor microenvironment (TME), an ecosystem consisting of cancer, immune, and stromal cells, is considered a critical factor influencing tumorigenesis, tumor spread, metastasis, and treatment response [Larson et al., 2025; Bilotta et al., 2022]. Systematic investigation of the TME is crucial for understanding the complex interplay between cancer cells and immune cells in their surrounding spatial regions, such as stromal, immune, and vascular components. The spatial organization of cells within the TME is fundamental to cancer biology, which drive key oncogenic processes such as tumor progression, immune evasion, and therapeutic resistance [Marusyk et al., 2020; Seferbekova et al., 2023]. Traditional bulk and single-cell sequencing techniques lack spatial information, limiting their ability to capture the complex architecture of the TME. In contrast, spatial transcriptomics (ST) has emerged as a transformative technology that bridges this gap by enabling high-resolution mapping of gene expression directly within spatial tissue architecture. These advances provide a powerful framework for characterizing tissue heterogeneity and for the study of key signaling pathways leading to novel spatial biomarker identifications [Dong and Zhang, 2022; Sun et al., 2020]. Modern ST technologies such as CosMx™ offer cellular resolution, and researchers often adopt a field-of-view (FOV)–based design, selecting representative spatial regions across distinct tumor areas. This approach enables high-resolution characterization of intra-tumor heterogeneity and spatial gradients within the TME.

### Multi-resolution ST in renal cancer

The motivating data for this work arise from a novel ST-based study in renal (kidney) cancer (see Figure 1 left panels). Specifically, a sarcomatoid renal cell carcinoma (sRCC) tissue obtained from a patient was profiled at multiple FOVs across the tumor using an imaging-based ST platform, CosMx™ Spatial Molecular Imager, to generate high throughput multi-resolution spatial gene expression (*n* = 960 genes) at the single-cell level (Figure 1A). At the “macro” level in Figure 1B, the FOVs capture a *tumor gradient* that was annotated by two genitourinary pathologists into three histopathological subtypes: sarcomatoid, clear cell, and transition. Sarcomatoid regions display aggressive, dedifferentiated phenotypes with loss of epithelial features and mesenchymal morphology [Delahunt et al., 2013], driven by epithelial–mesenchymal transition (EMT) programs [Boström et al., 2012; Čugura et al., 2024]. In contrast, clear cell regions retain epithelial characteristics and are generally less aggressive [May et al., 2024] with hypothesized transition regions being somehwere in between. At the “micro” cellular level within each of the (*n* = 23) FOVs (1C), single cells exhibit local niche specific gene expression dynamics (range: 399 to 1705; total 26K cells). This nested micro within macro architecture of tumors engenders the discovery of spatially varying regulatory biological mechanisms within local niches and global tumor gradients [Marusyk et al., 2020].

**Figure 1.**
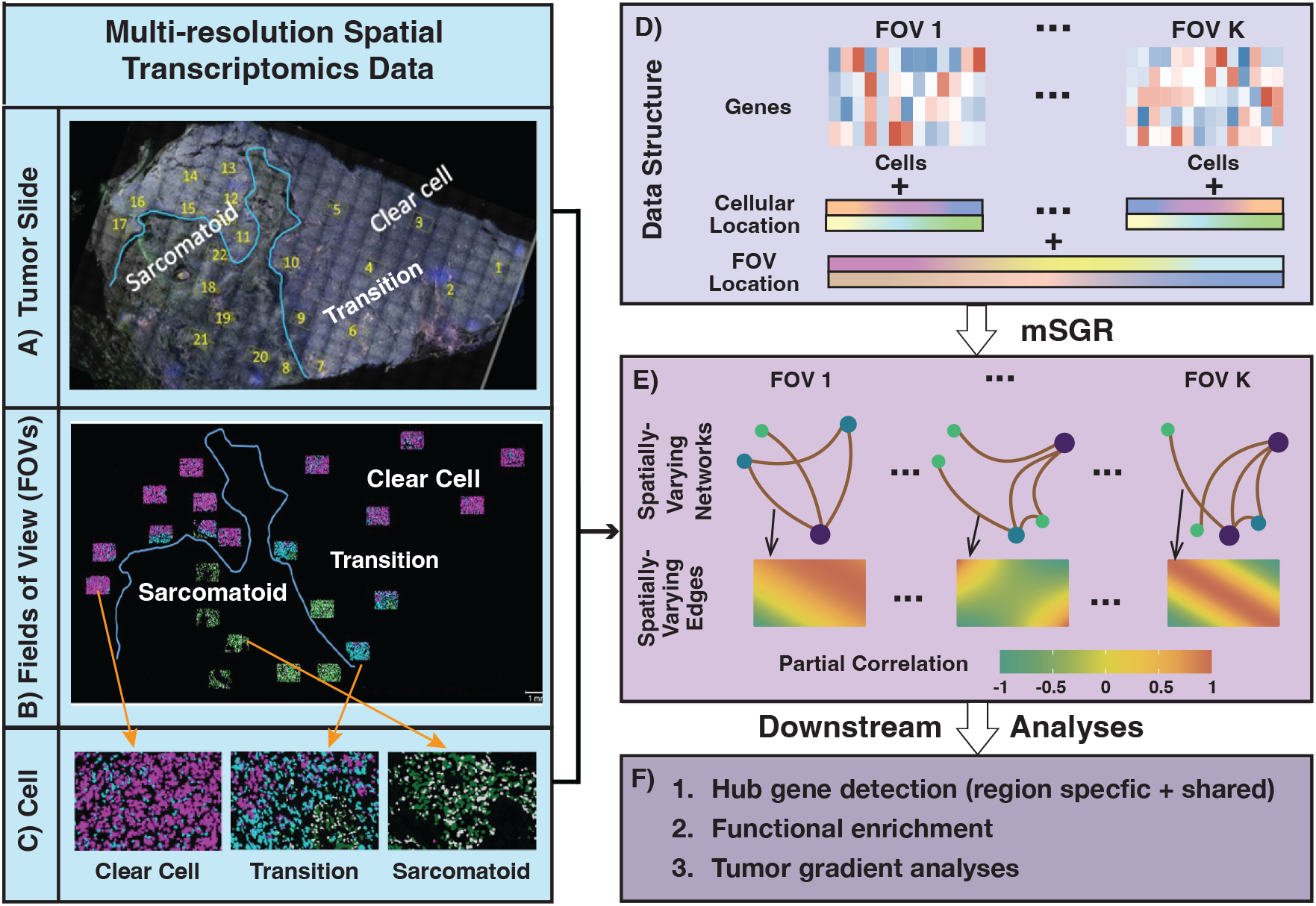
Overview of the Multi-resolution Spatial Graphical Regression (mSGR) framework: Tumor slide in Figure 1A is partitioned into multiple FOVs, each containing spatially resolved information on (i) FOV location within the tissue slide at a macro level (Figure 1B), and (ii) spatial coordinates of single cells at a micro level (Figure 1C). We implement the `mSGR` framework on hierarchical spatial gene expression data structure (Figure 1D) to infer multi-resolution spatially varying graphs to capture regional heterogeneity across the tissue (Figure 1E). In the downstream analyses on Figure 1F, we focus on hub gene identification (both region-specific and shared), functional enrichment of key pathways, and tumor gradient analyses across clear cell, transition, and sarcomatoid regions.

### Scientific problem

Several computational methods have been developed for ST data, primarily focusing on the identification of spatially variable genes [Sun et al., 2020; Yan et al., 2025] and spatial regions [Zhao et al., 2021; Dong and Zhang, 2022]. However, the inference of spatially varying genomic networks remains understudied. Genomic networks provide a system-level representation of molecular regulation, where nodes correspond to genes and edges encode some form of association or dependence. Modeling spatially varying gene networks enables the study of gene association patterns across different spatial regions in the TME. Network or graph-based perspectives are essential for studying regulatory mechanisms that are not captured by single-gene analyzes, thereby offering deeper insights into the organizational principles of spatial gene expression dynamics [Wang et al., 2025b].

In this paper, we aim to incorporate the spatial hierarchical data structure at both micro (within-FOV) and macro (between-FOV) scales (Figure 1D) and infer multi-resolution spatially varying graphs (Figure 1E). By leveraging spatial information from FOV locations along the tumor gradient, our framework provides a parsimonious representation of spatial network structures. These network summary measures can be used for downstream analyzes such as hub-gene identification and functional pathway enrichment (Figure 1F).

### Existing methods and limitations

Recent efforts have begun addressing spatial heterogeneity in network estimation for ST data. Neural-network–based algorithmic approaches integrate spatial adjacency or embedding information to infer gene–gene associations [Yuan and Bar-Joseph, 2020; Xu and McCord, 2021]. In contrast, modeling-based approaches represent spatial variation through probabilistic formulations of the underlying dependence structures. Broadly, these modeling-based methods can be classified into two categories in terms of modeling spatially varying marginal and conditional dependence through covariance and precision matrices, respectively. The marginal correlation modeling approaches estimate the joint structure through parametric or nonparametric covariance functions, often leveraging spatial kernels or decomposition techniques to account for spatial heterogeneity [Acharyya et al., 2022; Bernstein et al., 2022; Chakrabarti et al., 2024; Vasconcelos et al., 2025]. The conditional dependence–based approaches extend Gaussian graphical models (GGMs) to a spatial setting using spatially varying coefficient frameworks [Chen et al., 2025; Acharyya et al., 2025]. The conditional approaches yield sparser and biologically interpretable networks while mitigating the confounding effects from indirect associations. However, all of these methods are limited to analyzing a single spatial tissue domain or assume independence across samples in the case of multi-sample analyses.

In non-spatial settings, several approaches have been proposed for graphical model estimation for dependent samples, termed multiple GGMs (mGGMs), that jointly estimate a collection of precision matrices, enabling the characterization of shared and view specific conditional structures. Joint graphical lasso-based methods leverage structural similarity while accounting for group specific variation [Guo et al., 2011; Danaher et al., 2014]. Bayesian approaches for mGGMs link multiple graph structures through a Markov random field (MRF) prior [Peterson et al., 2015], spike and slab formulations [Li et al., 2019; Yang et al., 2021], or graphical horseshoe priors [Lingjærde et al., 2024; Busatto and Stingo, 2025] to accord network sparsity and heterogeneity. However, none of these methods incorporate spatial dependency or allow for the inference of spatially varying graphical structures. In summary, all of these aforementioned classes of methods are unable to infer spatially varying genomic graphs with a hierarchical multi-resolution spatial structure, as shown in Figure 1, incorporating both macro- and micro-level tumor information.

### Novel contributions

To address these challenges, we propose a multi-resolution spatial graphical regression (`mSGR`) – a Bayesian framework to infer spatially varying graphs across hierarchical spatial domains. Our model generalizes the mGGMs, conferring several key advantages. First, it flexibly models the precision matrix through Gaussian process priors, allowing the underlying graphical structure to vary across hierarchical spatial structures. Second, leveraging multi-resolution ST, the model borrows information across FOVs employing a Bayesian structured variable selection technique, where the edge-inclusion indicators incorporate the underlying topological spatial domain. This serves as a scalable alternative to MRF and structured spike-and-slab priors and allows for symmetry and positive definiteness constraints. Third, we impose a sparsity constraint to mimic the sparse nature of ST data, which enhances statistical interpretability and computational tractability. Finally, we propose a novel augmented variational inference algorithm to fit the `mSGR` model, which allows for parallelization and provides a scalable and efficient alternative to computationally intensive Markov Chain Monte Carlo (MCMC) methods for fitting such large-scale spatial datasets.

Comprehensive simulation studies demonstrate that the proposed `mSGR` framework accurately recovers spatially varying network structures across diverse settings. By jointly modeling multi-resolution spatial dependencies and spatially adaptive information sharing across FOVs, `mSGR` attains higher accuracy than many existing methods, underscoring its robustness and effectiveness in characterizing spatial heterogeneity. Analyses of sRCC ST data using `mSGR` reveal FOV-specific network structures of EMT genes, capturing spatially varying conditional dependencies between biologically linked gene pairs, such as collagen-encoding genes, and the dynamic hub behavior of key regulators such as MALAT1 and VIM along the tumor progression gradient. Furthermore, `mSGR` identifies markedly higher network connectivity in sarcomatoid regions, particularly within pathways associated with invasion, metastasis, and resistance to cell death, delineating the mechanistic rewiring accompanying aggressive tumor phenotypes.

### Sectional contents

The remainder of this paper is organized as follows. Section 2 introduces the proposed `mSGR` framework along with the model formulation, characterization, and prior specifications. The details of variational inference of `mSGR` are provided in Section 3. Section 4 presents comprehensive simulation studies to assess the performance of `mSGR` under different levels of spatial heterogeneity, data dimensions, and comparisons with competing methods. Section 5 applies `mSGR` to the multi-resolution ST from RCC tissue, where we study multi-resolution spatially varying graphs to infer spatial regulatory genes. We conclude with a discussion of methodological extensions and potential biomedical implications in Section 6. The `mSGR` R package is open-source and available on GitHub at https://github.com/***.

## 2 Multi-resolution spatial graphical models

We start by establishing the notational details for the multi-resolution ST data structure. Suppose a tissue slide is composed of *K* FOVs which are regions of interest where the *k*-th {*k ∈* (1, … , *K*)} FOV consists of *Nk* spatially indexed cells. The FOV specific spatial surface is denoted as 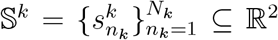(micro-resolution) and the whole spatial domain of the tissue slide can be represented as 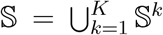 (macro-resolution). The observed gene expression matrix from *p* genes in the *k*th FOV is denoted as 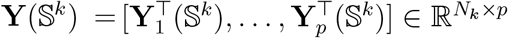.

Our inferential object is multi-resolution spatially varying graphs, which are characterized by *G*^*k*^[*V, E*^*k*^(𝕊^*k*^)]; where *V* denotes the vertex set of *p* genes and *E*^*k*^(𝕊^*k*^) denotes spatially varying edge set across *K* FOVs encoding spatially varying conditional dependencies. Our primary inferential goal is the spatially varying graph topology while borrowing information on *E*^*k*^(𝕊^*k*^) across FOVs. To elucidate the core construction, we first start with mGGM in Section 2 and generalize to spatial settings to introduce spatial mGGMs and `mSGR` in Section 2.1 and Section 2.2 respectively.

### Multiple Gaussian graphical models

Traditional GGMs encode conditional dependencies among variables through non-zero entries of the precision matrix [Dempster, 1972]. A standard (nonspatial) GGM is represented by an undirected graph *G* = (*V, E*), where edges represent non-zero partial correlations under a multivariate Gaussian model **Y** *∼ 𝒩* (0, **Ω**^*−*1^), with precision matrix **Ω**. Two nodes *i* and *j* are connected if their partial correlation *ρij* = Corr(**Y***i*, **Y***j* | **Y***l, l* ≠ *i, j*) is non-zero.

In mGMM settings, the observed data is sourced from *K* related views with 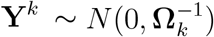 [Lingjærde et al., 2024]. For each *k*, the inferential object is a view-specific precision matrix 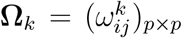, corresponding to a graph *G*^*k*^ = (*V, E*^*k*^). The goal is to jointly infer 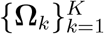, capturing both shared and view-specific graph structures. The *k*-th view specific partial correlation between node *i* and *j* is quantified as 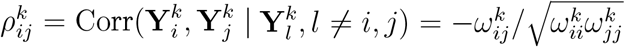 . This implies that 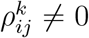 (or equivalently 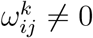) indicates conditional dependence between nodes *i* and *j* in the *k*-th graph. The graph structure can be represented with a collection of undirected graphs *G*^*k*^ = (*V, E*^*k*^), where *V* = {1, … , *p*}, 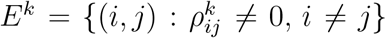 The joint graph estimation in mGGM setting reduces to identifying the non-zero elements of the precision matrices while borrowing strengths across the related views. However, these models impose a single shared graph for all cells within a given view i.e. assume independence between cells. Hence, we generalize mGGMs to accommodate spatial dependencies among cells within a given view (FOV in our case).

### 2.1 Multi-resolution spatially varying GGM

We introduce multiresolution spatial GGM which infers spatially varying genomic graphs while leveraging structured information across FOVs. We consider the joint distribution of **Y**(*s*^*k*^) = [*y*1(*s*^*k*^), … , *yp*(*s*^*k*^)] as

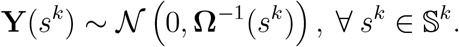

where **Ω**(*s*^*k*^) = {*ωij*(*s*^*k*^)}*p×p* is the spatially varying precision matrices across FOVs. Analogously to mGGM settings, the spatially varying partial correlation across FOVs is defined

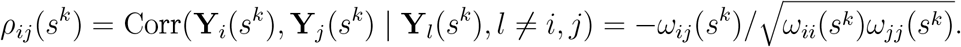

The corresponding multi-resolution spatial graph at location *s*^*k*^ is then denoted as, *G*^*k*^(*s*^*k*^) = [*V, E*^*k*^(*s*^*k*^) s.t. *E*^*k*^(*s*^*k*^) = *{*(*i, j*) : *i* ≠ *j*, s.t. *ω*_*ij*_(*s*^*k*^)≠ 0*}*], for any *s*^*k*^∈ 𝕊^*k*^.

Following the notions of mGGM, the conditional independence between two nodes given all other nodes at location *s*^*k*^ i.e. **Y**_*i*_(*s*^*k*^) ╨ **Y**_*j*_(*s*^*k*^) | **Y**_*l*_(*s*^*k*^) s.t. *l* ≠ {*i, j*} is noted by *ω*_*ij*_(*s*^*k*^) = 0. The conditional dependence of spatial graph *G*^*k*^(*s*^*k*^) structure can be characterized through non-zero elements of spatially varying precision matrices **Ω**(*s*^*k*^). The goal of this paper is to infer the non-zero elements of **Ω**(*s*^*k*^) while borrowing information across the *K* FOVs. Next, we introduce the `mSGR` framework to achieve our inferential goal in a parsimonious manner.

### 2.2 Multiresolution spatial graphical regression

The spatially varying precision matrices 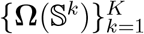 is a high dimensional object where the number of estimands are on the order of 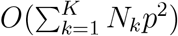. For example, for our case study where the number of cells are on the order of 10^4^ and even with moderate number of genes (*p* = 27 *∼* 115) this can exceed 10^6^ parameters – necessitating a computationally tractable estimation strategy. For example, a joint estimation strategy based on MRF [Peterson et al., 2015] or Wishart priors [Lenkoski and Dobra, 2011] becomes infeasible due computational complexity. To this end, we employ a neighborhood selection and regression-based approach [Meinshausen and Bühlmann, 2006; Ni et al., 2019; Ha et al., 2021; Zhang and Li, 2022] to estimate the non-zero elements of 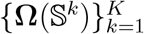.

Specifically, we introduce the `mSGR` model as

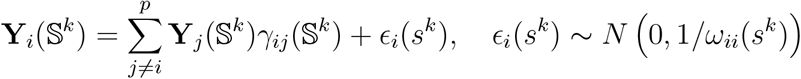

where *γ*_*ij*_(𝕊^*k*^) is interpreted as the spatial graphical regression coefficients between genes *i* and *j* within *k*th FOV. Since our primary focus is on estimating the off diagonal elements, we assume that diagonal variances *ω*_*ii*_(*s*^*k*^) does not vary with spatial location and can be represented by a constant value 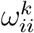. For any location *s*^*k*^ *∈ 𝕊*^*k*^ , *γ*_*ij*_(*s*^*k*^) is characterized as 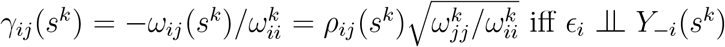. This establishes one-to-one correspondence between spatial graphical regression coefficients and conditional dependencies encoded in the spatially varying precision matrix Ω(*s*^*k*^).

To enable flexibility and sparsity in the estimation of the spatial genomics networks, we decouple the spatial graphical regression coefficients *γ*_*ij*_(𝕊^*k*^) as follows:

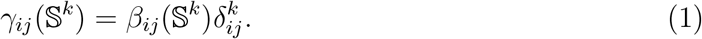

where *β*_*ij*_(𝕊^*k*^) is the magnitude of the spatial surface along with a selection indicator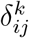.

#### Spatial surface modeling via Gaussian process

To admit a flexible spatial structure, we model *β*_*ij*_(𝕊^*k*^) with nonparametric Gaussian process prior with modified squared exponential kernel [Shi and Kang, 2015; Acharyya et al., 2025], enabling it to capture spatial variation within the FOVs. Using the Karhunen–Loève (KL) expansion, *β*_*ij*_(𝕊^*k*^) can be approximated by a set of basis functions *B*_*l*_(*·*) and corresponding coefficients 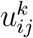, shown as:

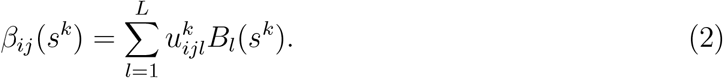

Further details regarding Gaussian process-based modeling of *β*_*ij*_(𝕊^*k*^) is deferred to Section S1.1 of the Supplementary Materials.

### 2.3 Bayesian structured edge selection across spatial surfaces

Our primary interest lies in modeling of selection indicator 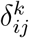, which represents the existence of a (spatial) edge between gene *i* and gene *j* in FOV *k*. Under the standard formulation, the indicators *δ* are assumed to be independent across spatial surfaces [Mitchell and Beauchamp, 1988]. However, in many applications, particularly those involving spatial, temporal, or network-structured data—selection, indicators often exhibit an underlying topographic or spatial structure, and the independence assumption is no longer hold [Andersen et al., 2017]. It is reasonable to expect that the joint inclusion of two selection indicators, *δ*_*i*_ and *δ*_*j*_, with spatial locations *s*_*i*_ and *s*_*j*_, depends on their proximity, such that ℙ (*δ*_*i*_ = 1, *δ*_*j*_ = 1) increases as |*s*_*i*_ *− s*_*j*_| decreases. This indicates variables with spa-tial proximity of each other are more likely to be jointly selected or excluded. This is particularly relevant in spatial omics, where genes co-expressed within the same tumor microenvironment often exhibit correlated regulation [Du et al., 2024; Wang et al., 2025a]. To accommodate such structured dependencies, several extensions of the spike-and-slab framework have been proposed. These include the use of logistic regression–based product priors to capture group-specific effects [Stingo et al., 2010; Menacher et al., 2024], Ising priors to encode spatial or graph-based similarities [Li and Zhang, 2010; Peterson et al., 2015], and structured spike-and-slab priors that introduce smooth spatial correlations among inclusion probabilities [Andersen et al., 2014, 2017].

Building on these ideas, we propose a novel Bayesian structured edge selection prior on the binary inclusion variables, following the framework of [Andersen et al., 2014; Mohammed et al., 2021]. Specifically, each indicator 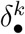 controls the inclusion of edge between nodes over the spatial domain 𝕊^*k*^ . These indicators are modeled as Bernoulli random variables with selection probabilities determined through a probit link,

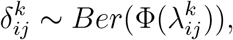

where Φ(*·*) denotes the standard normal cumulative distribution function. The latent vari-ables 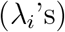 for each node-specific regression are collected into a matrix, and are assigned a matrix-variate Gaussian prior:

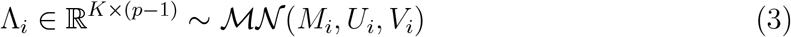

#### Characterization of *V*

The column covariance matrix *V*_*i*_ *∈ ℝ*^*K×K*^ captures the macro-level spatial correlations across FOVs. Each entry, denoted as *V*_*i*_[*k, k*^*′*^] reflects the prior correlation between FOVs *k* and *k*^*′*^, enabling the model to borrow strength across FOV level resolution. This structure aligns with multi-resolution ST, since neighboring FOVs are expected to exhibit similar biological processes. In our application, we parameterize *V*_*i*_ using a block-wise correlation structure. Specifically, FOVs belonging to different pathological regions (e.g., clear-cell, transitional, or sarcomatoid) are assumed to be independent, such that their cross-region correlations are set to zero. Within each region, the correlation between any two FOVs *k* and *k*^*′*^ is defined as

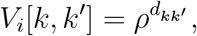

where 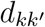 denotes the spatial distance between the centroids of FOVs *k* and *k*^*′*^, and *ρ ∈* (0, 1) controls the rate of spatial decay.

#### Characterization of *U*

The row covariance matrix *U*_*i*_ *∈* ℝ ^(*p−*1)*×*(*p−*1)^ encodes the dependencies among predictor genes while regressing on the *i*-th node. This covariance structure enables incorporation of prior knowledge of dependence between genes and coherent edge selection among biologically related genes in a pathway across spatial locations.

#### Characterization of *M*

The mean matrix is denoted by *M*_*i*_ *∈ ℝ*^*K×*(*p−*1)^, where each row corresponds to a specific field of view (FOV) and each column corresponds to one of the *p −* 1 potential predictor genes. This structure allows the model to incorporate prior knowledge about edge inclusion across spatial regions, and to capture both global and local regulatory signals.

In summary, the matrix normal prior Λ_*i*_ *∼ ℳ 𝒩* (*M*_*i*_, *U*_*i*_, *V*_*i*_) provides a structured and interpretable way to borrow strength and model prior beliefs about both spatial and genewise dependencies in the inclusion mechanism for edges in the graphical model. More specifically, let indicator being 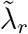denote the *r*-th element of vec(Λ_*i*_), with the corresponding selection 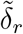. Under the matrix-variate Gaussian prior, the marginal distribution of 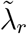 is Gaussian with mean 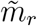 , the *r*-th element of vec(*M*_*i*_), and variance 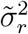, the *r*-th diagonal entry of *V*_*i*_ *⊗ U*_*i*_. From equations 3, the marginal prior distribution for 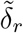 being selected is:

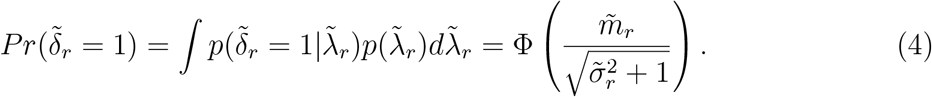

Equation 4 implies that the prior distribution for 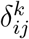 is non-informative when 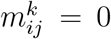 with the corresponding 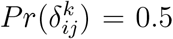, while 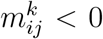 or 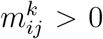 favor 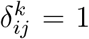 or 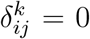 , respectively. Additionally, the marginal prior for 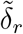 and 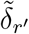 is simultaneously selected can be expressed as,

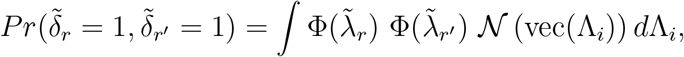

which captures the dependency in prior selection probabilities induced by the joint distribution of the latent variables and the ability of the model to account for correlation or shared structure in the inclusion. This is further exemplified in Figure 2A, which shows that higher prior means increase selection probabilities, while larger variances make this effect more gradual by reducing the influence of the mean. Similarly, a higher prior correlation among the latent Gaussian variables induces stronger concordance among the corresponding selection indicators (Figure 2B). In practice, this induced concordance is typically smaller than the specified prior correlation due to the nonlinear thresholding transformation from latent variables to binary indicators. Nevertheless, this mechanism effectively captures the spatial dependence among FOVs, as nearby FOVs with higher prior correlation are more likely to exhibit similar inclusion patterns. Consequently, the model encourages spatially coherent variable selection while allowing localized deviations supported by the data.

**Figure 2.**
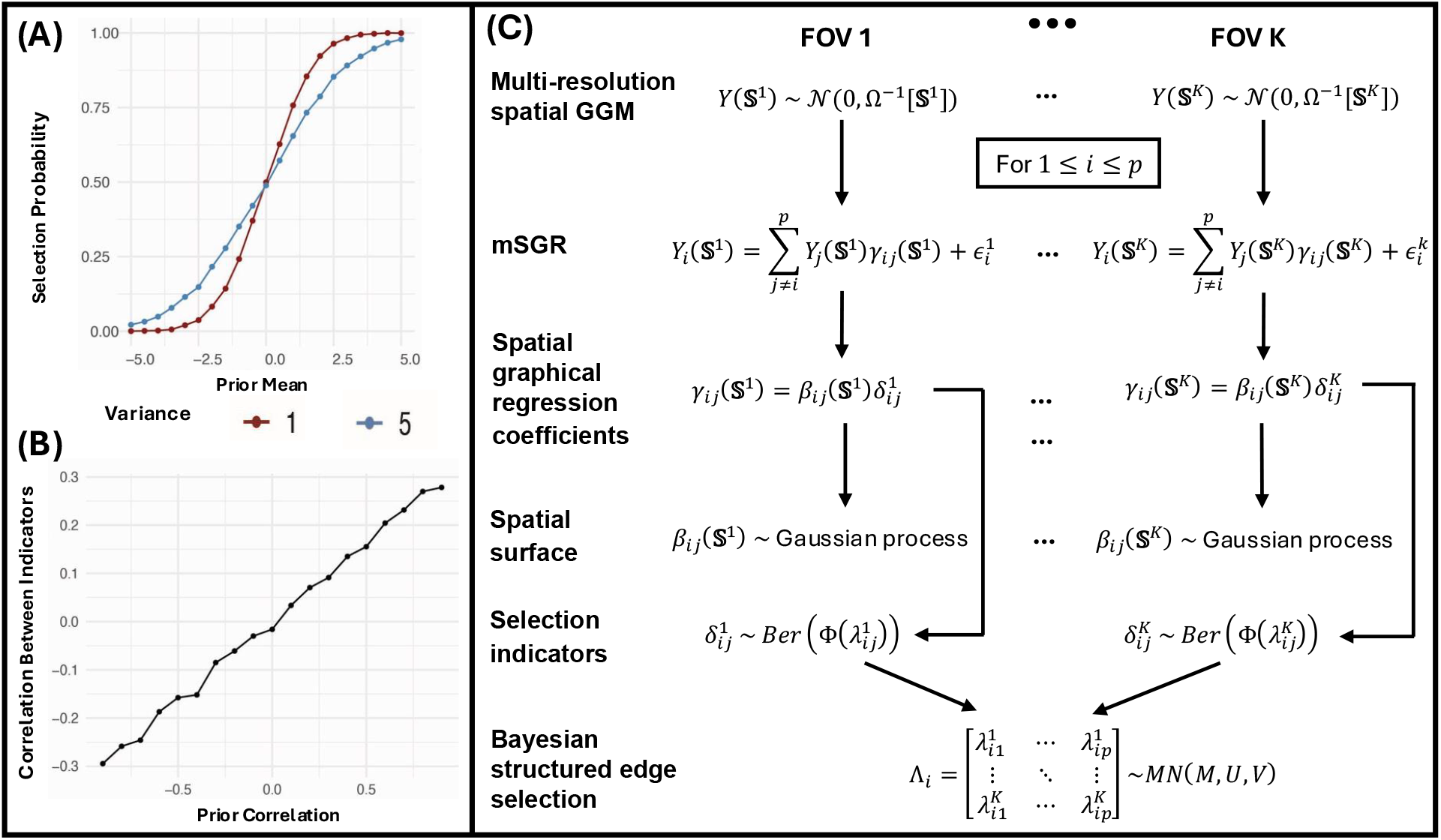
Overview of the Bayesian multiresolution spatial graphical regression (mSGR) model. **(A)** The impact of the prior mean on the marginal probability of selection for two variables with differing prior variances (1 in dark red, 5 in steel blue). As the prior mean increases, the probability of selection increases monotonically for both variables, with the higher-variance variable showing a more gradual change. **(B)** The influence of prior correlation on the empirical correlation between binary selection indicators. As the prior correlation increases, the resulting correlation between indicators also increases. **(C)** Flow diagram of the `mSGR` model.

#### mSGR model overview

Figure 2C summarizes the various components of the proposed `mSGR` model. In summary, for each FOV *k*, the conditional dependence among genes is modeled through spatially graphical regression, with coefficients 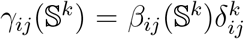. The continuous effects *βij*(𝕊^*k*^) follow Gaussian process priors to capture smooth spatial variation. The binary indicators 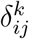 indicates the presence of an edge between genes *i* and *j* in FOV *k*. 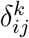 are assumed to follow Bernoulli 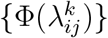, which are linked through a matrix-variate normal prior on latent variables Λ*i ∼* MN(*M, U, V*) to induce structured dependence across genes and FOVs. To complete prior specification, we assume a conjugate Gamma prior for the precision parameter, 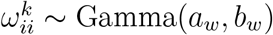.

## 3 Augmented variational inference for **mSGR**

Variational inference (VI) [Jordan et al., 1999] offers a scalable alternative to MCMC by reframing Bayesian inference as an optimization problem [Blei et al., 2017]. Instead of drawing samples from the posterior, VI approximates the posterior distribution through a target distribution by minimizing in-between Kullback–Leibler (KL) divergence. A standard choice of target distribution is the mean-field variational family [Wainwright et al., 2008], which assumes a fully factorized form over latent variables, thereby simplifying the optimization by decoupling the high-dimensional inference problem into a set of lower-dimensional ones. However, it also neglects dependencies among correlated variables, potentially leading to biased approximations [Blei et al., 2017]. Specifically, we define the structured variational distribution for the parameter space *θ* as

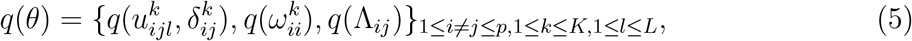

which allows for joint estimation of effect sizes 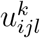 and their corresponding selection indicators 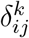. This formulation improves the approximation of posterior dependencies and provides uncertainty quantification, as it preserves the intrinsic coupling between the magnitude parameters and the corresponding selection indicators [Velten and Huber, 2021].

### Augmented Transformation

Since we place a probit prior on the selection indicators, i.e., 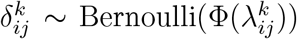 , the resulting model is non-conjugate. Conjugacy allows for analytical computation of variational expectations, thereby facilitating efficient and deterministic optimization of the evidence lower bound via coordinate-wise updates [Blei et al., 2017]. In the absence of conjugacy, these expectations often do not admit closed-form solutions. To address the intractability introduced by the probit prior, we use a data augmented framework for probit regression [Albert and Chib, 1993]. We introduce auxiliary latent variables 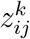 such that,

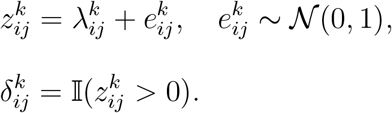

This augmentation transforms the non-conjugate probit model into a conditionally conjugate form, restoring tractability in variational updates. The binary selection indicators 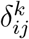 can be dealt with the continuous latent variables 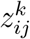 , which admit Gaussian conditional distributions. This reformulation enables efficient inference and retains compatibility with coordinate ascent updates within the structured MFVB framework (see Section S1.8).

### Symmetry and positive definiteness

To ensure the symmetry of the estimated conditional dependence matrices, i.e., *ωij*(*s*^*k*^) = *ωji*(*s*^*k*^) for each slide *k*, we adopt a post-processing strategy inspired by the symmetrization rules in [Meinshausen and Bühlmann, 2006]. Let the posterior inclusion probability (PIP) for the edge between nodes *i* and *j* in FOV *k* be defined as 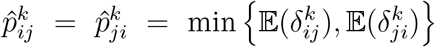. The conditional dependency exists, i.e., 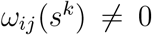, if 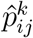 exceeds a threshold *κα*. To determine *κα*, we use a Bayesian local false discovery rate (FDR) control procedure that aims to control the average Bayesian FDR at a pre-specified level *α* [Baladandayuthapani et al., 2010]. For edges that satisfy 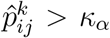, we define 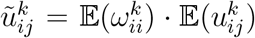. To enforce symmetry, we define the final symmetrized estimate by selecting the one with smaller *l*1 norm 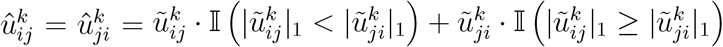 .

As the off-diagonal entries in the precision matrix are spatially varying, a natural sufficient condition for ensuring positive definiteness of Ω(*s*) is to make it diagonally dominant, *∀s ∈ S*. We implement a rescaling step in post processing procedure of 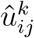 [Zhang and Li, 2022] to ensure 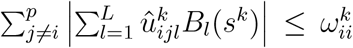 . Define *a*^*k*^ = ∥*B*^*k*^∥∞ < ∞, where ∥ · ∥∞ represents the sup-norm. Since *B*^*k*^ represents the basis functions in the Gaussian process described in Section S1.1, the quantity *a*^*k*^ is an observed value that depends on spatial location and the Gaussian process parameters. We then rescale the coefficients 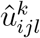 by the factor 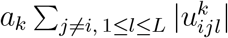 to satisfy the above condition. A detailed proof is provided in the Section S1.9.

## 4 Simulation Studies

To demonstrate the effectiveness of our proposed `mSGR` method, we conduct replicated simulations studies under various simulation settings to evaluate the performance in terms of graph structural recovery. We begin by outlining the data generation process, which mimics the multi-resolution architecture of spatial transcriptomics data, followed by descriptions of the competing methods and structural recovery measures. We assess the performance of `mSGR` under two scenarios: a synthetic setting under varying sample sizes and spatial correlations (Scenario I) and a sRCC data-informed simulation incorporating spatially varying edges (Scenario II). All results are aggregated over 50 independent replicates.

### Data generation for scenario I

We fix the number of genes at *p* = 30 with sparsity at 5% and vary the sample size for each FOV *Nk ∈* {500, 1000, 1500}. The spatial correlation matrix 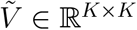 is defined with entries 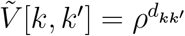, where 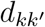 denotes the Euclidean distance between centroids of FOVs *k* and *k*^*′*^ and spatial correlation are ranged from *ρ ∈* {0.3, 0.6, 0.9}. FOVs are arranged in a 5 *×* 5 grid, each occupying a 0.4 *×* 0.4 unit square, with 0.6-unit spacing between adjacent regions. Within each FOV *k*, we generate a 40 *×* 40 regular grid of candidate cell positions and randomly sample *Nk* cells without replacement, yielding spatial coordinates 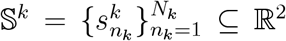. For each gene pair (*i, j*), Λ*ij ∈* ℝ^*K*^ is generated from a multivariate normal distribution i.e. 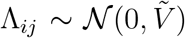, based on which selection indicators are defined using a threshold to control the sparsity. For each included edge, we consider a spatial function to quantify the off-diagonal entries and diagonal entries are adjusted to ensure positive definite property of precision matrices. The gene expression vectors are generated from the resulting multivariate Gaussian distributions.

### Data generation for scenario II

We adopt a similar structure from scenario I while mimicking real data based settings. The spatial coordinates and the number of cells within each FOV are considered from the RCC dataset in Section 5. We fix the number of genes at *p* = 27 and set the sparsity level to 12%, consistent with our observation from RCC data. To construct spatially varying edges, we construct a library of smooth functions derived from the estimated spatial effects 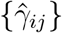 in RCC dataset. For each function, we compute its maximum absolute value across cells within FOVs and retain those with maximum magnitude exceeding 0.5, and scale them by a factor of 2. Each selected edge is randomly assigned to one of these rescaled functions to define the spatially varying off-diagonal elements in the precision matrix. To guarantee positive definiteness, we iteratively add a small quantity of 0.2 to all diagonal entries until the resulting matrix becomes positive definite. Finally, we simulate the gene expression vectors from multivariate Gaussian distributions with derived spatially varying precision matrices.

### Comparison Methods

We compare our proposed method against several state-of-the-art approaches for estimating multiple graphical models. Spatial Graphical Regression (SGR, Acharyya et al. [2025]) is a spatially aware method to estimate spatially varying graphs but it is limited to a single contiguous region and does not account for multiple FOVs. GraphR [Chen et al., 2025] is a probabilistic graphical regression framework that incorporates external covariates through a linear design matrix. We also compare `mSGR` with existing mGGM-based approaches. Joint Graphical Lasso (JGL, Danaher et al. [2014]) and jointGHS [Lingjærde et al., 2024] are multi-group Gaussian graphical models that estimate precision matrices for each FOV jointly with a group-fused penalty or hierarchical shrinkage, respectively. However, they assume discrete groups and do not incorporate spatial distances across regions. Finally, we include a baseline graphical lasso (Glasso, Friedman et al. [2008]) method applied independently to each FOV, which does not borrow information across FOVs and ignores spatial context entirely.

### Evaluation Metrics

For comprehensive assessment of graph structural recovery, we use three performance metrics: Matthews correlation coefficient (MCC), true positive rate (TPR), and false discovery rate (FDR). MCC incorporates all four components of confusion matrix components—providing a summary statistic reflecting structural recovery performance. Ranging from -1 to 1, an MCC of 1 indicates perfect classification, and -1 reflects total disagreement. In the context of sparse graph recovery, MCC is a robust metric for evaluating binary classification performance, especially in imbalanced settings [Matthews, 1975; Chicco and Jurman, 2020].

### Simulation results of scenario I

Figure 3A presents the simulation results under various degrees of correlation between FOVs and sample sizes. At a moderate correlation level (*ρ* = 0.6) and a sample size of 1000, `mSGR` achieves the highest value of MCC (`mSGR`: 0.9, SGR: 0.89, GraphR: 0.87, GLASSO: 0.72, GGL: 0.53, JointGHS: 0.32) among other competing methods. The `mSGR` method achieves moderately high TPR (`mSGR`: 0.84, SGR: 0.81, GraphR: 0.83, GLASSO: 0.8, GGL: 0.9, JointGHS: 0.11) while balancing for low FDR (`mSGR`: 0.021, SGR: 0.002, GraphR: 0.08, GLASSO: 0.314, GGL: 0.645, JointGHS: 0.002). GGL shows high TPR although it suffers from the highest FDR of 0.65 among all methods. SGR yields marginally lower FDR than `mSGR` though it comes at the cost of lower TPRs. `mSGR` achieves lower FDR than all other methods. In summary, `mSGR` consistently demonstrates balanced performance in terms of high precision, and reasonable control over false positives, outperforming other methods in MCC.

**Figure 3.**
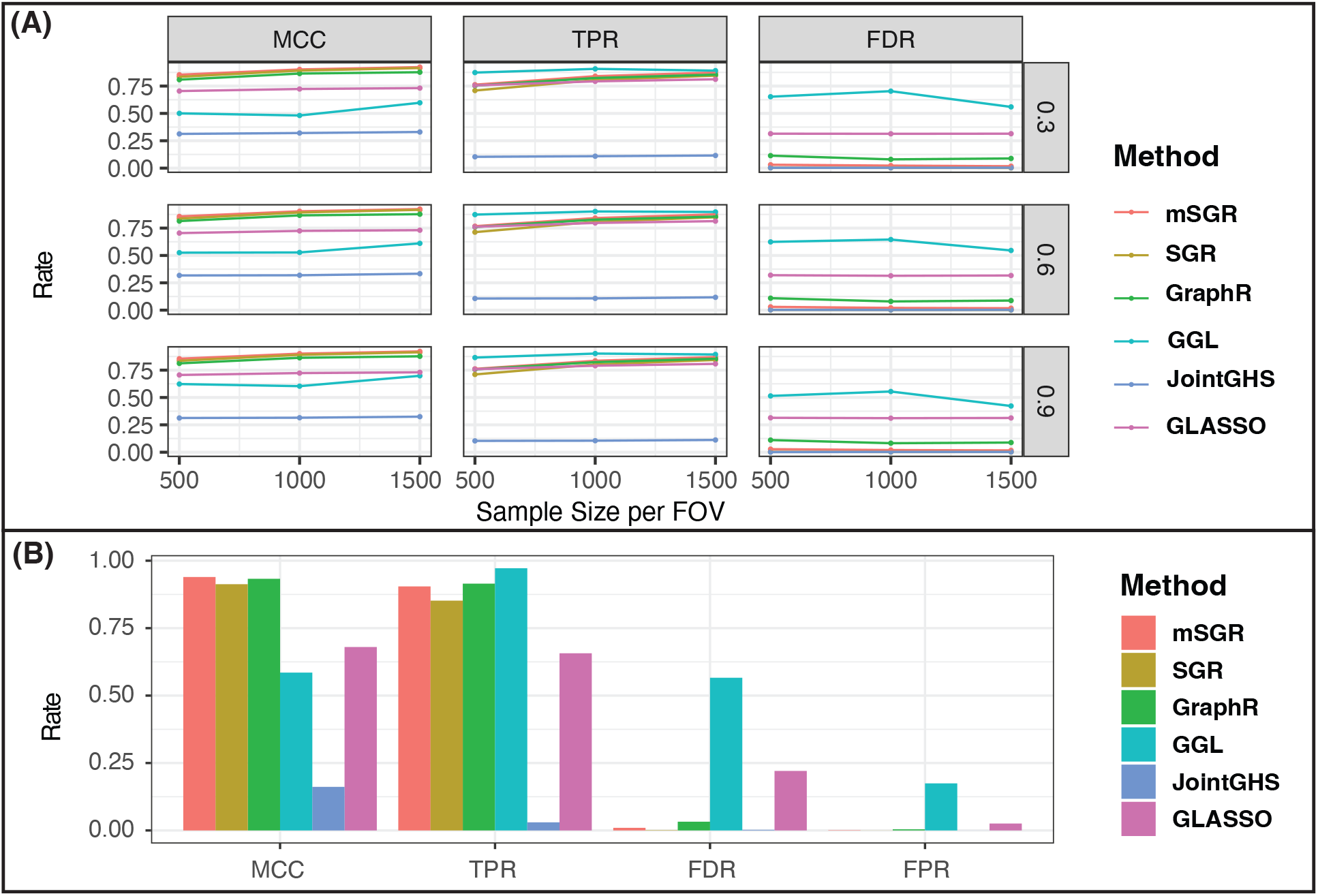
Simulation performance of mSGR. **(A)** Simulation results comparing different methods under varying correlation strengths between FOVs (*ρ* = 0.3, 0.6, 0.9) and sample sizes (*n* = 500, 1000, 1500 per FOV). Figure 3**(B)** shows competitive performance of `mSGR` for graph structral recovery in the real data–based simulation. The plot represents mean values of the evaluation metrics (MCC, TPR, FDR, FPR) across replications.

### Simulation results of scenario II

Figure 3B demonstrates the performance of `mSGR` under real data based simulation settings. The proposed `mSGR` (MCC: 0.94, TPR: 0.90, FPR: 0.001, FDR: 0.01) attains the highest MCC, reflecting a high level of agreement with the true underlying spatially varying graphs, along with a high TPR and very low error rates. The GraphR (MCC: 0.93, TPR: 0.91, FPR: 0.004, FDR: 0.03) method achieves lower MCC and marginally higher sensitivity, although at the cost of reduced precision, as indicated by higher levels of FDR and FPR. SGR (MCC: 0.91, TPR: 0.85, FPR < 0.001, FDR: 0.001) yields competitive results, with lower MCC than `mSGR` and very low FDR, FPR, but it shows relatively lower detection power with TPR. In contrast, the remaining methods exhibit performance limitations. GGL (MCC: 0.59, TPR: 0.97, FPR: 0.17, FDR: 0.57) achieves a high TPR, but its precision suffers significantly, as reflected through low FDR and MCC. GLASSO (MCC: 0.68, TPR: 0.66, FDR: 0.22, FPR: 0.03) yields a comparatively similar performance. The method jointGHS (MCC: 0.16, TPR: 0.03, FDR: 0.002, FPR: *≈* 0) provides very limited performance with overly conservative edge selection that fails to recover meaningful network structures despite minimal error rates.

### Computation time and additional simulation studies

In a high-performance computing environment, the average runtime per replicate for `mSGR` in Simulation I is approximately 0.34 hours with 500 samples per FOV, 1.26 hours with 1,000 samples per FOV, and 2.43 hours with 1,500 samples per FOV. In Simulation II, the average runtime per replicate is approximately 1.34 hours. For reference, running the application on a local machine (MacBook Pro equipped with an Apple M4 Pro chip) requires approximately 7 minutes. Further computational details are provided in the Section S2.4. We performed a detailed sensitivity analyses to assess the robustness of `mSGR` to prior specification and model flexibility. For different settings of Gaussian process hyperparameters and sparsity levels, `mSGR` shows consistent trends in MCC, TPR, FDR, and FPR as reported in the Section S2.3.

## 5 Multi-resolution spatial network characterization in renal cell carcinoma

### Scientific background

Renal cell carcinoma (RCC) is the most common type of kidney cancer [Cirillo et al., 2024]. In 2023, an estimated 81,800 new cases were diagnosed in the United States, making it the sixth most common cancer among males and ninth most common among females. Globally, RCC is the 15th leading cause of cancer-related death, with over 179,000 deaths reported in 2020 [Rose and Kim, 2024]. Sarcomatoid renal cell carcinoma (sRCC) is a de-differentiation of primary RCC which arises through epithelial-mesenchymal transition (EMT) [Čugura et al., 2024]. While sRCC represents the most extreme version of EMT in RCC, EMT plays a critical role in all RCC tumors broadly and is associated with an increased risk [Piva et al., 2016]. However, EMT is not a uniform process; rather, it occurs in concert with dynamic changes in the TME [Brabletz et al., 2021]. A key scientific question, therefore, is how EMT and its interactions with the TME are organized within tumor architecture. While EMT in RCC can be highly heterogeneous, sRCC represents an ideal model to study EMT gradients, as it exhibits a structured progression of EMT that can be detected histologically and interrogated at the microscale. Therefore, spatially characterizing EMT within the TME provides a unique opportunity to uncover tumor gradients that are obscured in bulk analyses, offering deeper insights into mechanisms of tumor aggressiveness and potential therapeutic vulnerabilities.

### Multi-resolution ST data

Our case study arises from high-resolution hierarchical ST data from RCC tissue using the CosMx platform [May et al., 2025]. The dataset includes 23 FOVs across 960 genes, capturing a total of 26,298 cancer cells, with cell counts per FOV ranging from approximately 460 to 1,800. These FOVs are categorized into three histopathological subtypes: 8 sarcomatoid, 12 clear cell, and 3 transitional regions exhibiting intermediate phenotypic characteristics between clear cell and sarcomatoid types. We investigate multiple biological pathways, with a primary focus on the EMT pathway (*p* = 27 genes), which plays a critical role in RCC progression, invasion, and therapeutic response. For our investigation, we focus on cancer cells to delineate how gene regulatory patterns change across the spatial EMT gradient. Further details of the data preprocessing steps and choice of model hyperparameters can be found in Supplementary Section S3.

### Spatially varying graphs across multiple FOVs

We apply the `mSGR` on RCC data, and Figure 4 shows spatially varying graphs of EMT pathway genes across different spatial domains of clear cell, transitional, and sarcomatoid regions (marked in different background colors). We observe particularly dense networks in the transitional region, as well as in clear cell, and sarcomatoid areas located near the transition zone (FOVs 8, 9, 20, and 23). While marginal correlations among mesenchymal genes are generally high in sarcomatoid regions due to the global upregulation of EMT programs [Boström et al., 2012], their conditional dependence structure is relatively weaker, as much of the co-expression appears to arise from shared associations with largely mediated through shared links with extracellular matrix remodeling and immune-interaction genes—such as *COL1A1*, rather than direct regulatory interactions [Nieto et al., 2016].

**Figure 4.**
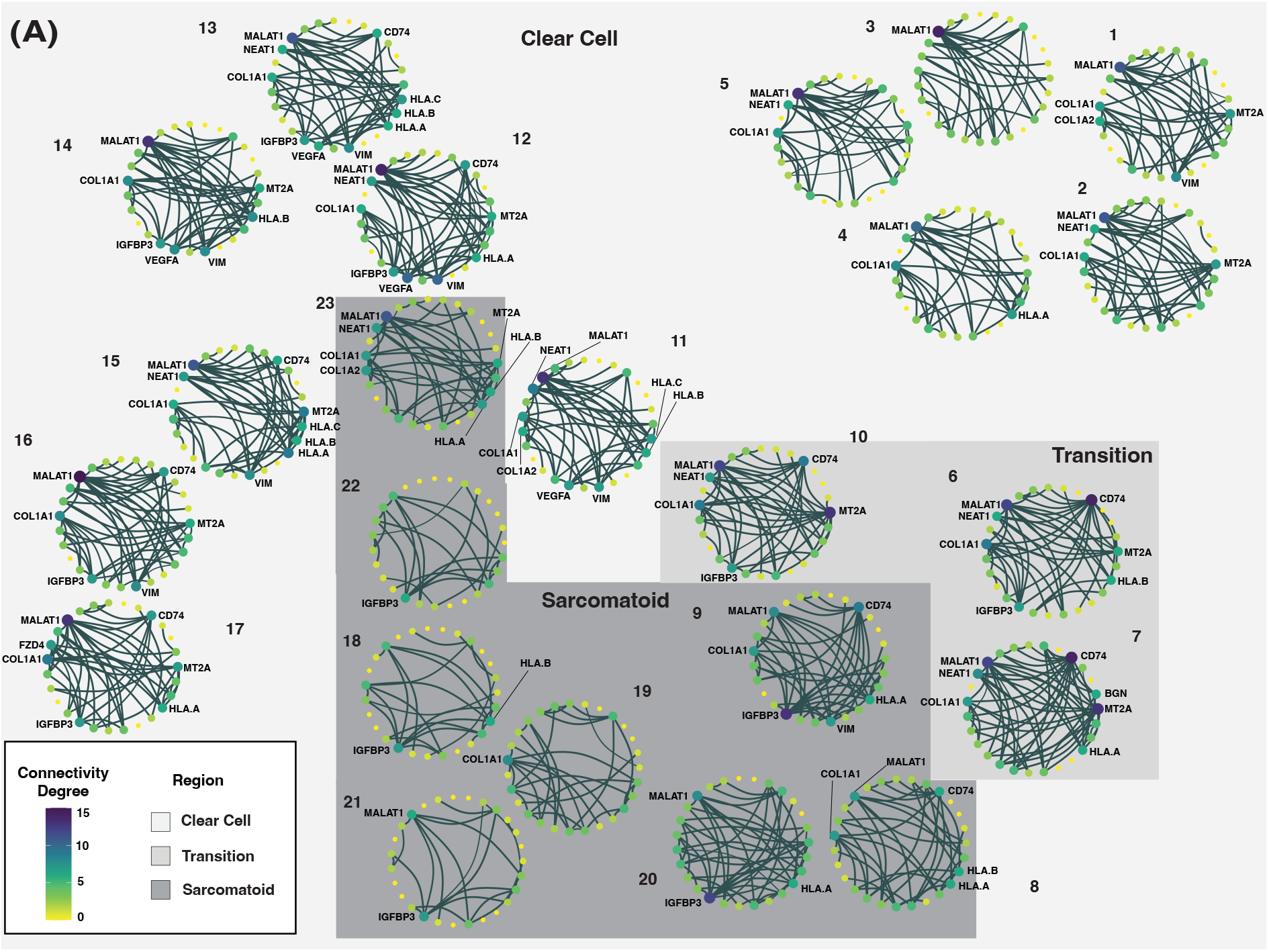
EMT Gene Networks Across Various Regions in RCC. (**A)** Networks depict conditional dependencies among genes from EMT pathway in clear cell, transitional, and sarcomatoid RCC regions. Nodes represent genes, and edges indicate conditional associations inferred from multi-resolution ST data. Node color corresponds to connectivity degree which is defined as the summation of significant connection, while posterior inclusion probabilities (PIP) quantify edge width. Gene names are displayed only for nodes with connectivity degree greater than 5. Each network is annotated with its corresponding FOV-specific number.

### Spatially varying edges

We present the spatially varying partial correlations of selected gene pairs in Figure 5A, demonstrating the ability of `mSGR` to incorporate spatial information and explicitly estimate heterogeneous dependence structures across regions. Notably, we observe a consistently high partial correlation between COL1A1 and COL1A2 across all regions, with the strongest signal emerging in the sarcomatoid area. This finding is biologically consistent, as the two genes encode the *α*1 and *α*2 chains of type I collagen, which physically assemble into a heterotrimeric triple helix [Li et al., 2016]. The statistical evidence of direct conditional dependence is concordant with their well-established structural interdependence in forming type I collagen. In the context of sarcomatoid, this tight coupling likely reflects enhanced extracellular matrix remodeling, which is closely associated with tumor progression and metastasis [Walker et al., 2018; Eble and Niland, 2019].

**Figure 5.**
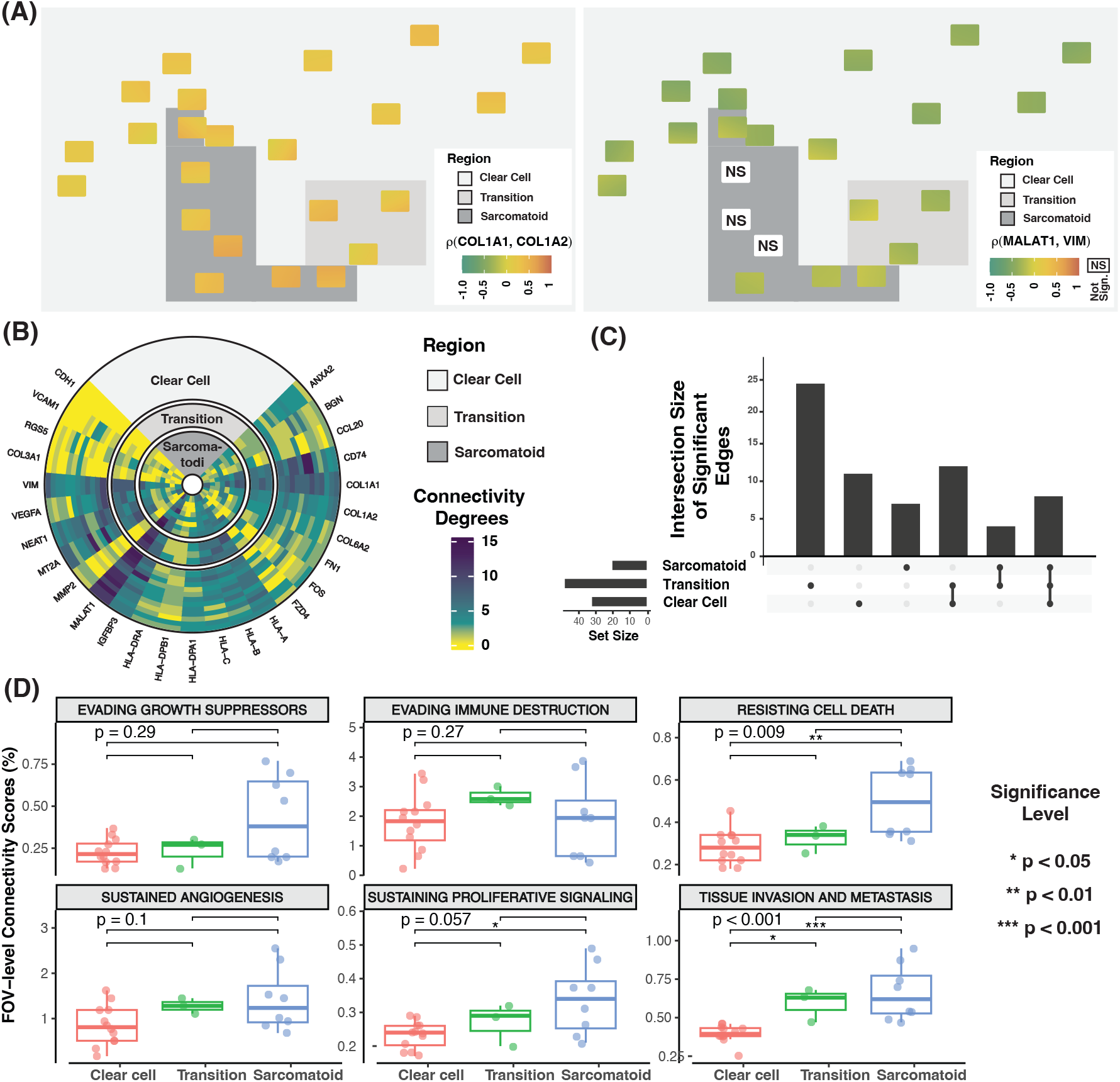
Genomic network-based characterization of region in RCC. (A) Spatial patterns of partial correlations for selected gene pairs (COL1A1–COL1A2; MALAT1–VIM), with background color distinguishing clear cell, transitional, and sarcomatoid regions. (B) Circular heatmap of connectivity degrees across genes within each region, where connectivity degree is defined as the total number of significant edges incident to a node. (C) UpSet plot of significant connections across regions, summarizing the number of significant gene–gene connections detected in each region and their overlaps. A connection is considered significant in a region if it is significant in more than 50% of the FOVs belonging to that region. (D) Boxplots showing the distribution of field-of-view (FOV)–level connectivity degree across spatial regions. The connectivity scores (CS) is defined as CS = # significant edges*/*# possible edges *×* 100. Each point represents one FOV, with regions color-coded. The overall difference among regions within each pathway facet was assessed using the Kruskal–Wallis test (p-values shown at the top), and pairwise differences were evaluated by Wilcoxon rank-sum tests with Benjamini–Hochberg adjustment. Significance levels are indicated by stars (* *p* < 0.05, ** *p* < 0.01, *** *p* < 0.001).

### Novel spatial biomarker identification

We also identify key hub genes by quantifying connectivity degrees, defined as the total number of significant edges connected to each node. Figure 5B highlights that MALAT1 and VIM consistently act as hub genes across all regions, exhibiting higher connectivity degrees in clear cell compared to sarcomatoid regions (also see Supplementary Figure S10). Additionally, to identify region-dependent hub patterns, we perform a nonparametric test to compare the connectivity degree across clear cell, transition, and sarcomatoid regions for all genes. Five genes (HLA-DPA1, MALAT1, MT2A, NEAT1 and VCAM1) show significant regional differences after multiple testing correction, indicating distinct patterns along the EMT gradient. Boxplots illustrating these differences are provided in Supplementary Figure S11.

Notably, MALAT1 and VIM show a strong negative partial correlation in clear cell regions, which becomes weaker in transition and sarcomatoid regions (Figure 5A right). MALAT1 is frequently upregulated in clear cell RCC and has been linked to poor prognosis [Zhang et al., 2015], although its role in sarcomatoid RCC remains less well characterized. In our data, MALAT1 expression is enriched in clear cell and transitional regions but decreases in sarcomatoid regions (Supplementary Figure S12) suggesting that its prognostic association may not directly translate to sarcomatoid differentiation. By contrast, VIM, a canonical mesenchymal marker—shows moderate expression in clear cell regions but strong enrichment in sarcomatoid regions [Čugura et al., 2024], consistent with its established role in mesenchymal activation. The reciprocal expression and negative conditional association between MALAT1 and VIM thus highlight an antagonistic relationship that evolves spatially along the tumor gradient, reflecting the dynamic balance between epithelial maintenance and mesenchymal activation during tumor progression.

Transitional regions exhibit stronger and more specific partial correlations, as illustrated in Figure 5C, where 47 significant edges (23 region-specific) are detected, compared with 31 (11 region-specific) in clear cell and 19 (7 region-specific) in sarcomatoid regions. Given that sarcomatoid differentiation has been strongly linked to EMT activation in RCC [Conant et al., 2011], the denser and more specific connectivity observed in transitional regions likely reflects dynamic regulatory rewiring during EMT progression, capturing the acquisition of mesenchymal features [Li et al., 2023; Pal et al., 2021]. These findings demonstrate that our framework captures spatial variation in network topology and identifies region-specific key regulators of RCC progression.

### Connectivity between genes in hallmark pathways

We next quantify the pathway coordination of 6 canonical hallmark pathways of cancer progression [Hanahan, 2022]. We use connectivity score (CS), defined as the proportion of significant partial correlations among all possible gene–gene pairs within a pathway. This metric captures the extant of coordinated molecular activity and provides a pathway-level summary of regulatory tightness [Chen et al., 2025]. Across regions, CS values reveal distinct functional remodeling along the clear cell–to–sarcomatoid transition (Figure 5D). These differences are statistically significant for both the resisting cell death (*p* = 0.009) and the tissue invasion and metastasis pathways (*p* < 0.001). For the resisting cell death pathway, the median CS increases from 0.28% (range: 0.18%-0.45%) in clear cell regions to 0.34% (range: 0.25%-0.38%) in transition regions and 0.5% (range: 0.31%-0.69%) in sarcomatoid regions. A similar pattern is observed for the tissue invasion and metastasis pathway, where sarcomatoid regions display consistently elevated connectivity (median 0.62%; range: 0.47%–0.95%) relative to clear cell regions (median 0.4%; range: 0.25%–0.46%). These observations align strongly with the known aggressive biology of sRCC, which is characterized by increased motility, invasive capacity, and a high resistance to apoptosis [Lebacle et al., 2019]. Moreover, sarcomatoid regions display a wider range of CS values across FOVs for most hallmark pathways, indicating greater heterogeneity in pathway-level network organization, consistent with previous findings that the mesenchymal component of sRCC exhibits a highly variable appearance [Lebacle et al., 2019].

## 6 Discussion

In this paper, we introduce the `mSGR` framework, a hierarchical Bayesian model for spatial network estimation of high-dimensional, and multi-resolution ST data. `mSGR` jointly infers multiple spatially varying graphs while borrowing information across spatial resolutions to capture both local heterogeneity and global spatial network structures. This is achieved through a novel Bayesian structured edge selection method, which regulates the inclusion probabilities of edges across spatial surfaces, allowing the graph topology to borrow strength from neighboring regions according both spatial distance and region annotations. The GP prior on spatially varying regression coefficients enables partial correlations to vary smoothly across spatial regions. This dual mechanism provides a coherent way to integrate global-local heterogeneity and multi-resolution spatial graphical inference.

Methodologically, `mSGR` generalizes existing spatial and multiple-GGM frameworks by embedding spatially structured Bayesian edge selection into the graphical model, and enabling spatially coherent yet locally adaptive edge selection. To achieve scalable and tractable inference, `mSGR` employs an MFVB algorithm built upon an augmentation representation that converts non-conjugate selection indicator priors into conditionally conjugate forms. This reparameterization allows closed-form variational updates, and enables computational efficiency while maintaining inference accuracy. Together, these innovations yield a unified, interpretable, and computationally efficient framework for analyzing spatially heterogeneous gene networks from multi-resolution designs.

Although we focus on undirected multiple spatially varying graphs, the `mSGR` framework can be adapted to more general classes of directed graphs with a known genomic ordering. Notably, this framework naturally extends to other continuous covariates, such as pathological gradients or clinical measures. From an inferential perspective, `mSGR` offers a principled way to quantify uncertainty in multiple spatially resolved network structures and to assess region-level connectivity metrics such as connectivity degrees and pathway-specific summaries. The hierarchical formulation also naturally accommodates multi-region comparisons and facilitates formal testing of region-dependent network alterations. While the motivating application focuses on ST data in cancer, the `mSGR` framework is broadly applicable to other domains involving spatially structured multivariate data, such as neuroimaging and environmental monitoring. Notably, `mSGR` also supports characterization of spatial networks across multiple (tissue) samples; in such settings, careful calibration of the sample-correlation prior *V* is important for robust borrowing of information across samples. Finally, `mSGR` can be extended to characterize networks for other spatial omics data and across spatial multi-omics layers; we leave these tasks for future consideration.

## SUPPLEMENTARY MATERIALS

### Title

The Supplementary PDF document provides additional methodological, computational and implementation details for `mSGR` along with additional results of simulation studies and ST data analysis.

### R-package

The R package is available at https://github.com/***.

### Data Availability Statement

Raw data of RCC are deposited in the Gene Expression Omnibus (GEO) under accession numbers GSE299368, and will be publicly available upon publication of the corresponding manuscript.

### Use of Generative AI

The authors used ChatGPT (OpenAI, GPT-5.2) to assist with language editing during manuscript preparation. The tool was used solely to refine grammar and improve readability and did not contribute to the methodology, analysis, or scientific conclusions. All authors have reviewed and approved the final manuscript.

## References

Acharyya, S., Kang, J., and Baladandayuthapani, V. (2025). Spatially varying graph estimation for spatial transcriptomics cancer data. bioRxiv, pages 2025–05.

Acharyya, S., Zhou, X., and Baladandayuthapani, V. (2022). Spacex: gene co-expression network estimation for spatial transcriptomics. Bioinformatics, 38(22):5033–5041.

Albert, J. H. and Chib, S. (1993). Bayesian analysis of binary and polychotomous response data. Journal of the American statistical Association, 88(422):669–679.

Andersen, M. R., Vehtari, A., Winther, O., and Hansen, L. K. (2017). Bayesian inference for spatio-temporal spike-and-slab priors. JMLR, 18(139):1–58.

Andersen, M. R., Winther, O., and Hansen, L. K. (2014). Bayesian inference for structured spike and slab priors. Advances in Neural Information Processing Systems, 27.

Baladandayuthapani, V., Ji, Y., Talluri, R., Nieto-Barajas, L. E., and Morris, J. S. (2010). Bayesian random segmentation models to identify shared copy number aberrations for array cgh data. Journal of the american statistical association, 105(492):1358–1375.

Bernstein, M. N., Ni, Z., Prasad, A., Brown, J., Mohanty, C., Stewart, R., Newton, M. A., and Kendziorski, C. (2022). Spatialcorr identifies gene sets with spatially varying correlation structure. Cell Reports Methods, 2(12).

Bilotta, M. T., Antignani, A., and Fitzgerald, D. J. (2022). Managing the tme to improve the efficacy of cancer therapy. Frontiers in Immunology, Volume 13-2022.

Blei, D. M., Kucukelbir, A., and McAuliffe, J. D. (2017). Variational inference: A review for statisticians. Journal of the American statistical Association, 112(518):859–877.

Boström, A.-K., Möller, C., Nilsson, E., Elfving, P., Axelson, H., and Johansson, M. E. (2012). Sarcomatoid conversion of clear cell renal cell carcinoma in relation to epithelial-to-mesenchymal transition. Human pathology, 43(5):708–719.

Brabletz, S., Schuhwerk, H., Brabletz, T., and Stemmler, M. P. (2021). Dynamic emt: a multi-tool for tumor progression. The EMBO journal, 40(18):e108647.

Busatto, C. and Stingo, F. C. (2025). Inference of Multiple High-Dimensional Networks with the Graphical Horseshoe Prior. JCGS, 0(0):1–12.

Chakrabarti, A., Ni, Y., and Mallick, B. K. (2024). Joint bayesian estimation of cell dependence and gene associations in spatially resolved transcriptomic data. Scientific Reports, 14(1):9516.

Chen, L., Acharyya, S., Luo, C., Ni, Y., and Baladandayuthapani, V. (2025). A probabilistic modeling framework for genomic networks incorporating sample heterogeneity. Cell Reports Methods, 5(2).

Chicco, D. and Jurman, G. (2020). The advantages of the matthews correlation coefficient over f1 score and accuracy in binary classification evaluation. BMC genomics, 21:1–13.

Cirillo, L., Innocenti, S., and Becherucci, F. (2024). Global epidemiology of kidney cancer. Nephrology Dialysis Transplantation, 39(6):920–928.

Conant, J. L., Peng, Z., Evans, M. F., Naud, S., and Cooper, K. (2011). Sarcomatoid renal cell carcinoma is an example of epithelial–mesenchymal transition. Journal of clinical pathology, 64(12):1088–1092.

Čugura, T., Boštjančič, E., Uhan, S., Hauptman, N., and Jeruc, J. (2024). Epithelial-mesenchymal transition associated markers in sarcomatoid transformation of clear cell renal cell carcinoma. Experimental and Molecular Pathology, 138:104909.

Danaher, P., Wang, P., and Witten, D. M. (2014). The joint graphical lasso for inverse covariance estimation across multiple classes. Journal of the Royal Statistical Society: Series B (Statistical Methodology), 76(2):373–397.

Delahunt, B., Cheville, J. C., Martignoni, G., Humphrey, P. A., Magi-Galluzzi, C., McKenney, J., Egevad, L., Algaba, F., Moch, H., Grignon, D. J., et al. (2013). The international society of urological pathology (isup) grading system for renal cell carcinoma and other prognostic parameters. The American journal of surgical pathology, 37(10):1490–1504.

Dempster, A. P. (1972). Covariance Selection. Biometrics, 28(1):157–175.

Dong, K. and Zhang, S. (2022). Deciphering spatial domains from spatially resolved transcriptomics with an adaptive graph attention auto-encoder. Nature communications, 13(1):1739.

Du, Y., Ding, X., and Ye, Y. (2024). The spatial multi-omics revolution in cancer therapy: Precision redefined. Cell Reports Medicine, 5(9).

Eble, J. A. and Niland, S. (2019). The extracellular matrix in tumor progression and metastasis. Clinical & experimental metastasis, 36(3):171–198.

Friedman, J., Hastie, T., and Tibshirani, R. (2008). Sparse inverse covariance estimation with the graphical lasso. Biostatistics, 9(3):432–441.

Guo, J., Levina, E., Michailidis, G., and Zhu, J. (2011). Joint estimation of multiple graphical models. Biometrika, 98(1):1–15.

Ha, M. J. et al. (2021). Bayesian structure learning in multilayered genomic networks. Journal of the American Statistical Association, 116(534):605–618. PMID: 34239216.

Hanahan, D. (2022). Hallmarks of Cancer: New Dimensions. Cancer Discovery, 12(1):31–46.

Jordan, M. I., Ghahramani, Z., Jaakkola, T. S., and Saul, L. K. (1999). An introduction to variational methods for graphical models. Machine learning, 37:183–233.

Larson, C. R., Mandloi, A., Acharyya, S., and Carstens, J. L. (2025). The tumor microenvironment across four dimensions: assessing space and time in cancer biology. Frontiers in Immunology, 16.

Lebacle, C., Pooli, A., Bessede, T., Irani, J., Pantuck, A. J., and Drakaki, A. (2019). Epidemiology, biology and treatment of sarcomatoid rcc: current state of the art. World journal of urology, 37(1):115–123.

Lenkoski, A. and Dobra, A. (2011). Computational Aspects Related to Inference in Gaussian Graphical Models With the G-Wishart Prior. JCGS, 20(1):140–157.

Li, D., Xia, L., Huang, P., Wang, Z., Guo, Q., Huang, C., Leng, W., and Qin, S. (2023). Heterogeneity and plasticity of epithelial–mesenchymal transition (emt) in cancer metastasis: Focusing on partial emt and regulatory mechanisms. Cell proliferation, 56(6):e13423.

Li, F. and Zhang, N. R. (2010). Bayesian variable selection in structured high-dimensional covariate spaces with applications in genomics. Journal of the American statistical association, 105(491):1202–1214.

Li, J., Ding, Y., and Li, A. (2016). Identification of col1a1 and col1a2 as candidate prognostic factors in gastric cancer. World journal of surgical oncology, 14(1):297.

Li, Z., Mccormick, T., and Clark, S. (2019). Bayesian joint spike-and-slab graphical lasso. Proceedings of the 36th International Conference on Machine Learning, 97:3877–3885.

Lingjærde, C., Fairfax, B. P., Richardson, S., and Ruffieux, H. (2024). Scalable multiple network inference with the joint graphical horseshoe. The Annals of Applied Statistics, 18(3):1899 – 1923.

Marusyk, A., Janiszewska, M., and Polyak, K. (2020). Intratumor heterogeneity: the rosetta stone of therapy resistance. Cancer cell, 37(4):471–484.

Matthews, B. W. (1975). Comparison of the predicted and observed secondary structure of t4 phage lysozyme. Biochimica et Biophysica Acta-Protein Structure, 405(2):442–451.

May, A., Williams, C., The, S., McGue, J., Wainstein, M., Shelley, G., Robinson, T., Salami, S. S., Mehra, R., Frankel, T., et al. (2024). Association of transitional gradient from clear cell to sarcomatoid renal cell carcinoma with macrophage/tumor cell crosstalk.

May, A. M., Kadomoto, S., Williams, C., Soupir, A. C., The, S., McGue, J. J., Robinson, T., Shelley, G., Hayes, M. T., Fridley, B. L., et al. (2025). Spatial analysis reveals a novel inflammatory tumor transition state which promotes a macrophage-driven induction of sarcomatoid renal cell carcinoma. bioRxiv, pages 2025–06.

Meinshausen, N. and Bühlmann, P. (2006). High-dimensional graphs and variable selection with the Lasso. The Annals of Statistics, 34(3):1436 – 1462.

Menacher, A., Nichols, T. E., Holmes, C., and Ganjgahi, H. (2024). Bayesian lesion estimation with a structured spike-and-slab prior. Journal of the American Statistical Association, 119(545):66–80.

Mitchell, T. J. and Beauchamp, J. J. (1988). Bayesian variable selection in linear regression. Journal of the american statistical association, 83(404):1023–1032.

Mohammed, S., Kurtek, S., Bharath, K., Rao, A., and Baladandayuthapani, V. (2021). Tumor radiogenomics with bayesian layered variable selection. arXiv preprint 2106.10941.

Ni, Y. et al. (2019). Bayesian graphical regression. Journal of the American Statistical Association, 114(525):184–197.

Nieto, M. A., Huang, R. Y.-J., Jackson, R. A., and Thiery, J. P. (2016). Emt: 2016. cell, 166(1):21–45.

Pal, A., Barrett, T. F., Paolini, R., Parikh, A., and Puram, S. V. (2021). Partial emt in head and neck cancer biology: a spectrum instead of a switch. Oncogene, 40(32):5049–5065.

Peterson, C., Stingo, F. C., and Vannucci, M. (2015). Bayesian inference of multiple gaussian graphical models. Journal of the American Statistical Association, 110(509):159–174.

Piva, F., Giulietti, M., Santoni, M., Occhipinti, G., Scarpelli, M., Lopez-Beltran, A., Cheng, L., Principato, G., and Montironi, R. (2016). Epithelial to mesenchymal transition in renal cell carcinoma: implications for cancer therapy. Molecular diagnosis & therapy, 20:111–117.

Rose, T. L. and Kim, W. Y. (2024). Renal cell carcinoma: A review. Jama, 332(12):1001– 1010.

Seferbekova, Z., Lomakin, A., Yates, L. R., and Gerstung, M. (2023). Spatial biology of cancer evolution. Nature Reviews Genetics, 24(5):295–313.

Shi, R. and Kang, J. (2015). Thresholded multiscale gaussian processes with application to bayesian feature selection for massive neuroimaging data. arXiv preprint 1504.06074.

Stingo, F. C., Chen, Y. A., Vannucci, M., Barrier, M., and Mirkes, P. E. (2010). A bayesian graphical modeling approach to microrna regulatory network inference. The annals of applied statistics, 4(4):2024.

Sun, S., Zhu, J., and Zhou, X. (2020). Statistical analysis of spatial expression patterns for spatially resolved transcriptomic studies. Nature methods, 17(2):193–200.

Vasconcelos, A. G., Danaher, P., McGuire, D., Wakefield, J., and Shojaie, A. (2025). Accounting for Spatial Correlation in Graphical Analysis of Spatial Transcriptomics Data. bioRxiv.

Velten, B. and Huber, W. (2021). Adaptive penalization in high-dimensional regression with external covariates using variational bayes. Biostatistics, 22(2):348–364.

Wainwright, M. J., Jordan, M. I., et al. (2008). Graphical models, exponential families, and variational inference. Foundations and Trends® in Machine Learning, 1(1–2):1–305.

Walker, C., Mojares, E., and del Río Hernández, A. (2018). Role of extracellular matrix in development and cancer progression. International journal of molecular sciences, 19(10):3028.

Wang, R., Qian, Y., Guo, X., Song, F., Xiong, Z., Cai, S., Bian, X., Wong, M. H., Cao, Q., Cheng, L., et al. (2025a). Stmodule: identifying tissue modules to uncover spatial components and characteristics of transcriptomic landscapes. Genome Medicine, 17(1):18.

Wang, X., Li, W. V., and Li, H. (2025b). spCLUE: a contrastive learning approach to unified spatial transcriptomics analysis across single-slice and multi-slice data. Genome Biology, 26(1):177.

Xu, Y. and McCord, R. P. (2021). CoSTA: unsupervised convolutional neural network learning for spatial transcriptomics analysis. BMC bioinformatics, 22(1):397.

Yan, G., Hua, S. H., and Li, J. J. (2025). Categorization of 34 computational methods to detect spatially variable genes from spatially resolved transcriptomics data. Nature Communications, 16(1):1141.

Yang, X., Gan, L., Narisetty, N. N., and Liang, F. (2021). GemBag: Group Estimation of Multiple Bayesian Graphical Models. JMLR, 22(54):1–48.

Yuan, Y. and Bar-Joseph, Z. (2020). GCNG: graph convolutional networks for inferring gene interaction from spatial transcriptomics data. Genome Biology, 21(1):300.

Zhang, H.-m., Yang, F.-q., Chen, S.-J., Che, J., and Zheng, J.-h. (2015). Upregulation of long non-coding rna malat1 correlates with tumor progression and poor prognosis in clear cell renal cell carcinoma. Tumor Biology, 36(4):2947–2955.

Zhang, J. and Li, Y. (2022). High-dimensional Gaussian Graphical Regression Models with Covariates. Journal of the American Statistical Association, 0(0):1–13.

Zhao, E., Stone, M. R., Ren, X., Guenthoer, J., Smythe, K. S., Pulliam, T., Williams, S. R., Uytingco, C. R., Taylor, S. E., Nghiem, P., et al. (2021). Spatial transcriptomics at subspot resolution with bayesspace. Nature biotechnology, 39(11):1375–1384.

