## Supplementary Material for "Multi-resolution Spatial Graphical Regression Models for Hierarchical Spatial Transcriptomics Data" (from another version of this preprint: version 2, 2026-05-16)

May 13, 2026

This Supplementary Material provides additional details to support the methodology, computation, and empirical results. Section S1 presents the `mSGR` model formulation and key methodological components, including the Gaussian process prior with the modified squared exponential kernel, model specification, priors and hyperparameters, initialization, the likelihood, derivation of closed-form MFVB updates, the full variational inference algorithm, and postprocessing steps such as Bayesian FDR thresholding and positive-definiteness enforcement. Section S2 provides additional simulation study details, including evaluation metrics, competing methods, and sensitivity analyses. Finally, Section S3 gives supplementary details for the real-data application on EMT gradient characterization in sarcomatoid renal cancer, including preprocessing/normalization, incorporation of spatial coordinates at both FOV and cell levels, hyperparameter choices, and additional figures.

#### S1 Methodological details for `mSGR`

##### S1.1 Gaussian process prior with modified exponential sqaured covariance kernel

We utilize Gaussian processes (GPs) to model spatial correlations on micro (cell) level. GPs serve as a fundamental tool for modeling complex spatial dependencies (Cressie and Wikle, 2011; Banerjee et al., 2003). In this context, we consider a zero-mean GP prior defined over a spatial domain $\mathbb{S}^k \subseteq \mathbb{R}^2$, characterized by a covariance kernel $\mathbf{C}(\cdot,\cdot) : \mathbb{R}^{2\times 2} \to \mathbb{R}$, such that $\beta(\cdot) \sim \mathcal{GP}(\mathbf{0}, \sigma^2\mathbf{C}(\cdot,\cdot))$. Under this formulation, any finite collection of spatial locations $s^k, t^k \in \mathbb{S}^k$ yields random variables that are jointly Gaussian distributed with covariance given by $\sigma^2\mathbf{C}(s^k, t^k)$.

In this work, we adopt a GP prior with a Modified Squared Exponential (MSE) kernel

(Shi and Kang, 2015; Lin et al., 2024) defined as:

$$\begin{aligned}&\beta_{ij}(\cdot) \sim \mathcal{GP}(\mathbf{0}, \sigma_{ij}^2 \mathbf{C}(\cdot,\cdot)\\&\mathbf{C}(s^k, t^k) = \exp\left\{-a_{gp}(||s^k||_2^2 + ||t^k||_2^2) - b_{gp}||s^k - t^k||_2^2\right\},\\&\text{for prespecified } a_{gp}, b_{gp} > 0, \text{ and } s^k, t^k \in \mathbb{S}^k.\end{aligned} \tag{S1}$$

Here $\|\cdot\|_2$ denotes the Euclidean norm. This kernel generalizes the classical squared exponential kernel by introducing an additional exponential decay factor dependent on the norms of $s^k$ and $t^k$. From the specified Gaussian process, the following properties are easily derived:

$$\begin{aligned}&\text{Var}[\beta_{ij}(0)] = \sigma_{ij}^2\\&\text{Var}[\beta_{ij}(s^k)] = \sigma_{ij}^2 \exp(-a_{gp}\|s^k\|_2^2)\\&\text{Cor}[\beta_{ij}(s^k), \beta_{ij}(t^k)] = \exp(-b_{gp}\|s^k - t^k\|_2^2)\end{aligned} \tag{S2}$$

Specifically, $\sigma_{ij}^2$ controls the maximum marginal variance of the process, setting the overall scale of variability for $\beta_{ij}(\cdot)$. The term $\exp(-a_{gp}(||s^k||_2^2 + ||t^k||_2^2)$ imposes global attenuation that diminishes influence far from the origin, with larger $a_{gp}$ corresponds to a faster decay rate of $\text{Var}[\beta_{ij}(s^k)]$ to $\text{Var}[\beta_{ij}(0)]$. Notably, when $a_{gp} = 0$, the modified squared exponential kernel becomes a standard exponential kernel. Meanwhile, the term $\exp(-b_{gp}||s^k - t^k||_2^2)$ captures local smoothness and spatial proximity, with a larger $b_{gp}$ corresponding to a smoother $\beta_{ij}(s^k)$, and thus impose stronger correlation between nearby points.

The primary barrier to implementing GPs lies in their significant computational and numerical challenges. For each field of view (FOV), GP inference requires inversion of an $N_k \times N_k$ covariance matrix, incurring a computational complexity of $O(N_k^3)$, which becomes prohibitively expensive as the dataset size grows. In addition to the computational cost, the inversion of the covariance matrix is numerically unstable, particularly when the matrix is ill-conditioned due to near-collinearity in the inputs or unfavorable kernel parameter settings (Banerjee et al., 2013). To facilitate computation and representation, we utilize Mercer's theorem (Minh et al., 2006; Shi and Kang, 2015), which states that under mild regularity conditions, a continuous, symmetric, and positive semi-definite kernel admits a spectral decomposition in terms of its eigenfunctions and eigenvalues. This result enables the application of the Karhunen–Loève (KL) expansion (Wang, 2008), expressed as

$$\beta_{ij}(s^k) = \sum_{l=1}^{\infty} u_{ijl}^k \sqrt{\eta_l}\psi_l(s^k) := \sum_{l=1}^{\infty} u_{ijl}^k B_l(s^k) \tag{S3}$$

where $\eta_l$ and $\psi_l(\cdot)$ are the eigenvalue-eigenfunction pairs of the covariance kernel, and

$u_{ijl}^k \overset{\text{i.i.d.}}{\sim} \mathcal{N}(0, \sigma_{ij}^2)$. This representation provides a compact and flexible approximation of the underlying stochastic process. In practical implementations, the infinite series is truncated to the first $L$ dominant components, thereby reducing the dimensionality from $N_k$ to $L$ and enabling scalable inference in high-dimensional spatial models.

#### S1.2 Augmented Model for Selection Indicators

As shown in Equation 3, we introduce a latent variable $z_{ij}^k = \lambda_{ij}^k + \epsilon_{ij}^k$, where $\epsilon_{ij}^k \sim N(0, 1)$. The selection indicator is then expressed as $\delta_{ij}^k = I(z_{ij}^k > 0)$. Consequently,
$P(\delta_{ijk} = 1) = P(z_{ij}^k > 0) = P(\lambda_{ij}^k + \epsilon_{ijk} > 0) = P(\epsilon_{ijk} > -\lambda_{ij}^k) = \Phi(\lambda_{ij}^k)$, where $\Phi(\cdot)$ denotes the cumulative distribution function of the standard normal distribution.

#### S1.3 Mean Field Variation Bayes

Suppose $\theta = \left\{u_{ijl}^k, \delta_{ij}^k, \omega_{ii}^k, \Lambda_{ij}\right\}_{1 \leq i \neq j \leq p, 1 \leq k \leq K, 1 \leq l \leq L}$ denote the parameters of interest. The true posterior distribution is $p(\theta \mid \mathcal{D})$, where $\mathcal{D}$ is the observed data. The goal is to approximate $p(\theta \mid \mathcal{D})$ with a structured variational distribution $q(\theta) \in \mathcal{Q}$, where $\mathcal{Q}$ includes factorizations that retain dependencies within predefined variable blocks, e.g.,

$$q(\theta) = q(u_{ijl}^k, \delta_{ij}^k) q(\omega_{ii}^k) q(\Lambda_{ij}) \tag{S4}$$

The optimal variational distribution $q^*(\theta)$ minimizes the KL divergence to the true posterior:

$$q^*(\theta) = \arg\min_{q \in \mathcal{Q}} \mathrm{KL}\left(q(\theta) \,\|\, p(\theta \mid \mathcal{D})\right).$$

This is equivalent to maximizing the evidence lower bound (ELBO):

$$\mathcal{L}(q) = \mathbb{E}_{q(\theta)}[\log p(\mathcal{D}, \theta)] - \mathbb{E}_{q(\theta)}[\log q(\theta)]$$

To improve numerical stability, all variational updates are computed using the full dataset. At iteration $t$, each variational parameter is updated according to

$$\boldsymbol{\theta}^{(t)} = \rho\, \boldsymbol{\theta}_{\text{new}}^{(t)} + (1 - \rho)\, \boldsymbol{\theta}^{(t-1)}, \tag{S5}$$

where $\rho = 0.9$ is the learning rate, $\boldsymbol{\theta}_{\text{new}}^{(t)}$ denotes the MFVB update obtained from the current coordinate ascent step, and $\boldsymbol{\theta}^{(t-1)}$ is the estimate from the previous iteration. This under-relaxation scheme improves convergence behavior and reduces oscillations in high-dimensional settings.

#### S1.4 Model, Priors, and Hyperparameters

For completeness, we summarize the model specifications, prior distributions, and associated hyperparameters used in the posterior computation. For each gene $i$ and FOV $k$, conditional on $\{u^k_{ijl}, \delta^k_{ij}, \omega^k_{ii}\}$, the observation vector $Y_i(\mathbb{S}^k)$ follows

$$Y_i(\mathbb{S}^k) \mid \{u^k_{ijl}, \delta^k_{ij}, \omega^k_{ii}\} \sim \mathcal{N}\left(\sum_{j\neq i} Y_j(\mathbb{S}^k) \otimes \gamma_{ij}(\mathbb{S}^k)\, I(z^k_{ij} > 0),\ (\omega^k_{ii})^{-1} I_{N_k}\right).$$

**Residual precision:** $\omega^k_{ii} \sim \text{Gamma}(a_\omega, b_\omega)$.

**GP coefficients (KL-based):** For $\tilde{v}^k_{ijl} = u^k_{ijl}\sqrt{\eta_l}, \tilde{v}^k_{ijl} \sim N(0,\ \sigma^2_{gp}\eta_l)$.

**Edge-selection latent variables (probit):**

$$z^k_{ij} = \lambda^k_{ij} + \epsilon^k_{ij}, \qquad \epsilon^k_{ij} \sim N(0,1),$$
$$\delta^k_{ij} = I(z^k_{ij} > 0), \qquad P(\delta^k_{ij} = 1) = \Phi(\lambda^k_{ij}).$$

**Spatial proximity prior on structured parameters:** $\Lambda_{ij} \sim \mathcal{N}(m_{\Lambda i,\text{prior}},\ \sigma^2_{\Lambda i} U)$.

**Hyperparameters:** Throughout, the hyperparameters $a_\omega,\ b_\omega,\ \sigma^2_{gp},\ m_{\Lambda i,\text{prior}},\ \sigma^2_{\Lambda i},\ U$ are treated as known and fixed. In simulation studies, we set $a_w = 10, b_w = 10, m_{\Lambda i,\text{prior}}, = 0, \sigma^{-2}_{\Lambda i} = 50$. For GP parameters, we set $\sigma^2_{gp} = 1, a_{gp} = 0.01, b_{gp} = 0.5$, approximation degree $= 10$, leading to $L = 66$.

**Notational details.** For field of view (FOV) $k$ with $N_k$ spatial locations and a truncated basis expansion of size $L$, we define the following quantities.

1. **Design matrix.** For each pair $(i, j)$, let

$$H^k_{ij} \in \mathbb{R}^{N_k \times L},$$

   where the $l^{\text{th}}$ column is given by

$$\left(H^k_{ij}\right)_{\cdot l} = Y_j(\mathbb{S}^k) \otimes \psi_l(\mathbb{S}^k).$$

2. **Scaled basis coefficients.** Define the scaled coefficients

$$\tilde{v}^k_{ijl} := u^k_{ijl}\sqrt{\eta_l}, \qquad \tilde{v}^k_{ijl} \sim \mathcal{N}(0, \eta_l),$$

for $l = 1, \ldots, L$.

3. **Coefficient vector.** Collect the scaled coefficients into the vector

$$\tilde{v}_{ij}^k = \left(\tilde{v}_{ij1}^k, \ldots, \tilde{v}_{ijL}^k\right)^\top \in \mathbb{R}^L.$$

4. **Basis expansion of spatial interaction.** The spatially varying regression term admits the representation

$$Y_j(\mathbb{S}^k) \otimes \gamma_{ij}(\mathbb{S}^k) = \sum_{l=1}^{L} \left(Y_j(\mathbb{S}^k) \otimes \psi_l(\mathbb{S}^k)\right) \tilde{v}_{ijl}^k = H_{ij}^k \tilde{v}_{ij}^k \in \mathbb{R}^{N_k}.$$

5. **Partial residual.** Define the partial residual excluding the $(i, j)$ interaction as

$$R_{ij}^k = Y_i(\mathbb{S}^k) - \sum_{j' \neq i,j} \left\{Y_{j'}(\mathbb{S}^k) \otimes \gamma_{ij'}(\mathbb{S}^k)\right\} I\left(z_{ij'}^k > 0\right).$$

6. **Quadratic form for the prior.** The Gaussian prior penalty can be written as

$$\sum_{l=1}^{L} \frac{(\tilde{v}_{ijl}^k)^2}{2\sigma_{ijk}^2 \eta_l} = \frac{1}{2\sigma_{ijk}^2} (\tilde{v}_{ij}^k)^\top \operatorname{diag}\left(\frac{1}{\eta_1}, \ldots, \frac{1}{\eta_L}\right) \tilde{v}_{ij}^k,$$

where we denote

$$\hat{\eta}^{-1} := \operatorname{diag}\left(\frac{1}{\eta_1}, \ldots, \frac{1}{\eta_L}\right).$$

#### S1.5 Initialization of Spatial Coefficients

We initialize the random variables $u_{ijl}^k$ in a field-of-view (FOV)–specific manner. For each FOV $k = 1, \ldots, K$, we extract the corresponding subsets of the response vector $\mathbf{Y}_i(\mathbb{S}^k)$, the predictor matrix $\mathbf{Y}_{-i}(\mathbb{S}^k)$, and the spatial basis matrix $B(\mathbb{S}^k) := \left[B_1(\mathbb{S}^k) \cdots B_L(\mathbb{S}^k)\right]$. A joint design matrix is then constructed by element-wise multiplying each predictor $\mathbf{Y}_{-i}(\mathbb{S}^k)$ with the spatial basis functions, yielding a matrix of dimension $n_k \times \{L(p-1)\}$. Using this joint design matrix, we fit a normal linear model via mean-field variational Bayes to obtain posterior mean estimates of the regression coefficients. These estimates are reshaped into an $L \times (p-1)$ matrix and used as the initial values for the coefficient matrix $u_{ij}^k$.

#### S1.6 Likelihood

$$
\begin{aligned}
L =& \prod_{i=1}^{p} \prod_{k=1}^{k} \left\{ \det([\omega_{ii}^{k}]^{-1} I_{N_k})^{-\frac{1}{2}} \exp\left( -\frac{\omega_{ii}^{k}}{2} \| Y_i(\mathbb{S}^k) - \sum_{j \neq i}^{p} Y_j(\mathbb{S}^k) \otimes \gamma_{ij}(\mathbb{S}^k) \cdot I(z_{ij}^{k} > 0) \|^2 \right) \right. \\
& \left. (\omega_{ii}^{k})^{a_\omega - 1} \exp(-b_\omega \cdot \omega_{ii}^{k}) \right\} \\
& \prod_{i=1}^{p} \prod_{j \neq i}^{p} \prod_{k=1}^{k} \prod_{l=1}^{L} \left\{ (\sigma_{gp}^{2} \eta^{l})^{-\frac{1}{2}} \exp\left( -\frac{(\tilde{v}_{ijl}^{k})^2}{2\sigma_{gp}^{2} \eta^{l}} \right) \right\} \\
& \prod_{i=1}^{p} \prod_{j \neq i}^{p} \prod_{k=1}^{k} \left\{ \exp\left( -\frac{1}{2} (z_{ij}^{k} - \lambda_{ij}^{k})^2 \right) \right\} \\
& \prod_{i=1}^{p} \prod_{j \neq i}^{p} \left\{ \exp\left( -\frac{1}{2} (\Lambda_{ij} - m_{\Lambda i,\text{prior}})^T (\sigma_{\Lambda i}^{2} U)^{-1} (\Lambda_{ij} - m_{\Lambda i,\text{prior}}) \right) \right\}.
\end{aligned}
$$

#### S1.7 Augmented Mean Field Variational Bayes (MFVB)

1. Update $\omega_{ii}^{k}$:

$$
\omega_{ii}^{k} \sim \text{Gamma}\left( \frac{N_k}{2} + a_\omega, \frac{1}{2} \mathbb{E} \left\| Y_i(\mathbb{S}^k) - \sum_{j \neq i} H_{ij}^{k} \cdot \tilde{v}_{ij}^{k} \cdot I(z_{ij}^{k} > 0) \right\|^2 + b_\omega \right).
$$

2. Update $\Lambda_{ij}$:

$$
\begin{aligned}
\mathbb{E}_{-\Lambda_{ij}} l =& \mathbb{E}_{-\Lambda_{ij}} \left\{ -\frac{1}{2} (\Lambda_{ij} - Z_{ij})^T (\Lambda_{ij} - Z_{ij}) - \right. \\
& \left. \frac{1}{2\sigma_{\Lambda i}^{2}} (\Lambda_{ij} - m_{\Lambda i,\text{prior}})^T U^{-1} (\Lambda_{ij} - m_{\Lambda i,\text{prior}}) \right\}
\end{aligned}
$$

$$
\Lambda_{ij} \sim \mathcal{N}\left( \left( I + \frac{1}{\sigma_{\Lambda i}^{2}} U^{-1} \right)^{-1} \left( \mathbb{E} Z_{ij} + \frac{1}{\sigma_{\Lambda i}^{2}} U^{-1} m_{\Lambda i,\text{prior}} \right), \left( I + \frac{1}{\sigma_{\Lambda i}^{2}} U^{-1} \right)^{-1} \right).
$$

3. Update $\tilde{v}_{ij}^k \mid z_{ij}^k$:

$$\begin{aligned}\log q(\tilde{v}_{ij}^k, z_{ij}^k) = \mathbb{E}_{-(\tilde{v}_{ij}^k, z_{ij}^k)}\Bigg[ &- \frac{\omega_{ii}^k}{2}\Big\| Y_i(\mathbb{S}^k) - \sum_{j\neq i} H_{ij}^k \tilde{v}_{ij}^k I(z_{ij}^k > 0)\Big\|^2 \\ &- \frac{1}{2}(z_{ij}^k - \lambda_{ij}^k)^2 - \frac{1}{2\sigma_{gp}^2}(\tilde{v}_{ij}^k)^T \hat{\eta}^{-1}\tilde{v}_{ij}^k\Bigg],\end{aligned}$$

$$\begin{aligned}\log q(\tilde{v}_{ij}^k \mid z_{ij}^k) &= \mathbb{E}_{-\tilde{v}_{ij}^k \mid z_{ij}^k}\Bigg[ - \frac{\omega_{ii}^k}{2}\Big\| Y_i(\mathbb{S}^k) - \sum_{j'\neq i} H_{ij'}^k \tilde{v}_{ij'}^k I(z_{ij'}^k > 0)\Big\|^2 \\ &\qquad - \frac{1}{2\sigma_{gp}^2}(\tilde{v}_{ij}^k)^T \hat{\eta}^{-1}\tilde{v}_{ij}^k\Bigg], \\ &= \mathbb{E}_{-\tilde{v}_{ij}^k \mid z_{ij}^k}\Bigg[ - \frac{\omega_{ii}^k}{2}\Big\| R_{ij}^k - H_{ij}^k \tilde{v}_{ij}^k I(z_{ij}^k > 0)\Big\|^2 - \frac{1}{2\sigma_{gp}^2}(\tilde{v}_{ij}^k)^T \hat{\eta}^{-1}\tilde{v}_{ij}^k\Bigg].\end{aligned}$$

Thus,

$$\tilde{v}_{ij}^k \mid z_{ij}^k \sim \mathcal{MVN}\Big(\hat{\mu}_{\tilde{v}}(z_{ij}^k), \hat{\Sigma}_{\tilde{v}}(z_{ij}^k)\Big).$$

with

$$\begin{aligned}\hat{\Sigma}_{\tilde{v}}(z_{ij}^k) &= \left[\mathbb{E}\omega_{ii}^k I(z_{ij}^k > 0)\|H_{ij}^k\|^2 + \mathbb{E}\frac{1}{\sigma_{gp}^2}\hat{\eta}^{-1}\right]^{-1}, \\ \hat{\mu}_{\tilde{v}}(z_{ij}^k) &= \hat{\Sigma}_{\tilde{v}}(z_{ij}^k)\cdot \mathbb{E}\omega_{ii}^k\left[I(z_{ij}^k > 0)\right](H_{ij}^k)^T \mathbb{E}R_{ij}^k.\end{aligned}$$

4. Update $z_{ij}^k$:

$$q(\tilde{v}_{ij}^k, z_{ij}^k)$$
$$= \exp\left\{\mathbb{E}\left[-\frac{\omega_{ii}^k}{2}\|R_{ij}^k - H_{ij}^k\tilde{v}_{ij}^k \cdot I(z_{ij}^k > 0)\|^2 - \frac{1}{2\sigma_{gp}^2}(\tilde{v}_{ij}^k)^T\hat{\eta}^{-1}\tilde{v}_{ij}^k - \frac{1}{2}(z_{ij}^k - \lambda_{ij}^k)^2\right]\right\}$$

$$q(z_{ij}^k) = \int q(\tilde{v}_{ij}^k, z_{ij}^k)d\tilde{v}_{ij}^k$$
$$= \det(\hat{\Sigma}_{\tilde{v}}(z_{ij}^k))^{1/2}\exp\left\{\frac{1}{2}\hat{\mu}_{\tilde{v}}^T(z_{ij}^k)\hat{\Sigma}_{\tilde{v}}^{-1}(z_{ij}^k)\hat{\mu}_{\tilde{v}}(z_{ij}^k)\right\}\exp\left\{-\frac{1}{2}(z_{ij}^k)^2 + \mathbb{E}\lambda_{ij}^k z_{ij}^k\right\}$$
$$\sim p_{ij}^k \cdot TN(\mathbb{E}\lambda_{ij}^k, 1, 0, +\infty) + (1 - p_{ij}^k) \cdot TN(\mathbb{E}\lambda_{ij}^k, 1, -\infty, 0)$$

$$p_{ij}^k = q(z_{ij}^k > 0) = \int_0^{+\infty} q(z_{ij}^k)dz_{ij}^k$$
$$= \det(\hat{\Sigma}_{\tilde{v}}(z_{ij}^k > 0))^{1/2} \cdot \exp\left\{\frac{1}{2}\hat{\mu}_{\tilde{v}}^T(z_{ij}^k > 0)\hat{\Sigma}_{\tilde{v}}^{-1}(z_{ij}^k > 0)\hat{\mu}_{\tilde{v}}(z_{ij}^k > 0)\right\}$$
$$\Phi(\mathbb{E}\lambda_{ij}^k) \cdot \text{Const}$$

$$1 - p_{ij}^k = q(z_{ij}^k < 0) = \int_{-\infty}^0 q(z_{ij}^k)dz_{ij}^k$$
$$= \det(\hat{\Sigma}_{\tilde{v}}(z_{ij}^k < 0))^{1/2} \cdot [1 - \Phi(\mathbb{E}\lambda_{ij}^k)] \cdot \text{Const}$$

$$\text{logit}(p_{ij}^k) = \frac{1}{2}\left[\log\det(\hat{\Sigma}_{\tilde{v}}(z_{ij}^k > 0)) - \log\det(\hat{\Sigma}_{\tilde{v}}(z_{ij}^k < 0))\right] +$$
$$\frac{1}{2}\hat{\mu}_{\tilde{v}}^T(z_{ij}^k > 0)\hat{\Sigma}_{\tilde{v}}^{-1}(z_{ij}^k > 0)\hat{\mu}_{\tilde{v}}(z_{ij}^k > 0) + \text{logit}(\Phi(\mathbb{E}\lambda_{ij}^k)).$$

**Numerical note:** When $\mathbb{E}[\lambda_{ij}^k]$ is positively or negatively large (e.g., $|\mathbb{E}[\lambda_{ij}^k]| > 10$), we have $\Phi(\mathbb{E}[\lambda_{ij}^k]) \to 1$ or 0, and hence $\text{logit}(\Phi(\mathbb{E}[\lambda_{ij}^k]))$ is not well-defined numerically. Using Mills' ratio $\lim_{x\to\infty} \frac{1-\Phi(x)}{\frac{1}{\sqrt{2\pi}}e^{-x^2/2}} = \frac{1}{x}$, we obtain the following approximations:

1. When $x$ is positively large, $1 - \Phi(x) \approx \frac{1}{\sqrt{2\pi}} \cdot \frac{e^{-x^2/2}}{x}$. Thus

$$\log\left(\frac{\Phi(x)}{1 - \Phi(x)}\right) \approx \frac{1}{2}\log 2\pi + \log x + \frac{x^2}{2}.$$

2. When $x$ is negatively large, $\Phi(x) \approx -\frac{1}{\sqrt{2\pi}} \cdot \frac{e^{-x^2/2}}{x}$, and

$$\log\left(\frac{\Phi(x)}{1-\Phi(x)}\right) \approx -\frac{1}{2}\log 2\pi - \log(-x) - \frac{x^2}{2}.$$

**Expression of expected values:**

1. $\mathbb{E}\|Y_i(\mathbb{S}^k) - \sum_{j\neq i}^{p} H_{ij}^k \tilde{v}_{ij}^k \cdot I(z_{ij}^k > 0)\|^2$

   Denote $G_{ijk} = \mathbb{E}(H_{ij}^k \tilde{v}_{ij}^k \cdot I(z_{ij}^k > 0)) = p_{ij}^k H_{ij}^k \hat{\mu}_{\tilde{v}}(z_{ij}^k > 0)$ and $\tilde{G}_{ik} = \sum_{j\neq i}^{p} G_{ijk}$. Then

$$\begin{aligned}
&\mathbb{E}\|Y_i(\mathbb{S}^k) - \sum_{j\neq i}^{p} H_{ij}^k \tilde{v}_{ij}^k \cdot I(z_{ij}^k > 0)\|^2 \\
=&\|Y_i(\mathbb{S}^k)\|^2 + \sum_{j\neq i}^{p} \mathbb{E}\|H_{ij}^k \tilde{v}_{ij}^k \cdot I(z_{ij}^k > 0)\|^2 - 2Y_{ik}^T(S)\mathbb{E}\left[\sum_{j\neq i}^{p} H_{ij}^k \tilde{v}_{ij}^k \cdot I(z_{ij}^k > 0)\right] + \\
&\sum_{j\neq i}^{p}\sum_{j'\neq i,j}^{p} \mathbb{E}\left[I(z_{ij}^k > 0)(\tilde{v}_{ij}^k)^T (H_{ij}^k)^T H_{ij'}k \tilde{v}_{ij'}^k I(z_{ij'k} > 0)\right].
\end{aligned}$$

   (a) $\mathbb{E}\|H_{ij}^k \tilde{v}_{ij}^k \cdot I(z_{ij}^k > 0)\|^2$:

$$\begin{aligned}
&\mathbb{E}\|H_{ij}^k \tilde{v}_{ij}^k \cdot I(z_{ij}^k > 0)\|^2 \\
=&p_{ij}^k \left[\hat{\mu}_{\tilde{v}}^T(z_{ij}^k > 0)([H_{ij}^k]^T H_{ij}^k)\hat{\mu}_{\tilde{v}}(z_{ij}^k > 0) + \mathrm{tr}\left([H_{ij}^k]^T H_{ij}^k \hat{\Sigma}_{\tilde{v}}(z_{ij}^k > 0)\right)\right] \\
=&\frac{1}{p_{ij}^k} G_{ijk}^T G_{ijk} + p_{ij}^k \cdot \mathrm{tr}\left([H_{ij}^k]^T H_{ij}^k \hat{\Sigma}_{\tilde{v}}(z_{ij}^k > 0)\right).
\end{aligned}$$

   (b) $Y_{ik}^T(S) \cdot \mathbb{E}\left[\sum_{j\neq i} H_{ij}^k \tilde{v}_{ij}^k \cdot I(z_{ij}^k > 0)\right] = Y_{ik}^T(S) \cdot \tilde{G}_{ik}$.

   (c) $\sum_{j\neq i}^{p}\sum_{j'\neq i,j}^{p} \mathbb{E}\left[I(z_{ij}^k > 0)(u_{ij}^k)^T (H_{ij}^k)^T H_{ij'k} u_{ij'k} I(z_{ij'k} > 0)\right]$:

$$\begin{aligned}
&\sum_{j\neq i}^{p}\sum_{j'\neq i,j}^{p} \mathbb{E}\left[I(z_{ij}^k > 0)(u_{ij}^k)^T (H_{ij}^k)^T H_{ij'k} u_{ij'k} I(z_{ij'k} > 0)\right] \\
=&\sum_{j\neq i}^{p}\sum_{j'\neq i,j}^{p} \left[G_{ijk}^T G_{ij'k}\right] = \sum_{j\neq i}^{p}\left[G_{ijk}^T(\tilde{G}_{ik} - G_{ijk})\right] = \tilde{G}_{ik}^T \tilde{G}_{ik} - \sum_{j\neq i}^{p} G_{ijk}^T G_{ijk}.
\end{aligned}$$

2. $\mathbb{E}[(\tilde{v}_{ij}^k)^T \tilde{v}_{ij}^k]$:

$$\begin{aligned}
&\mathbb{E}[(\tilde{v}_{ij}^k)^T \tilde{v}_{ij}^k] \\
=&\mathbb{E}_{z_{ij}^k}\left\{\mathbb{E}[(\tilde{v}_{ij}^k)^T \tilde{v}_{ij}^k | z_{ij}^k)]\right\} \\
=&\mathbb{E}_{z_{ij}^k}\left[\mathbb{E}^T(\tilde{v}_{ij}^k \mid z_{ij}^k)\mathbb{E}(\tilde{v}_{ij}^k \mid z_{ij}^k) + \mathrm{tr}(\mathrm{Var}(\tilde{v}_{ij}^k \mid z_{ij}^k))\right] \\
=&p_{ij}^k \cdot \left[\hat{\mu}_{\tilde{v}}(z_{ij}^k > 0)^T \hat{\mu}_{\tilde{v}}(z_{ij}^k > 0) + \mathrm{tr}(\hat{\Sigma}_{\tilde{v}}(z_{ij}^k > 0))\right] + (1 - p_{ij}^k)\mathrm{tr}(\hat{\Sigma}_{\tilde{v}}(z_{ij}^k < 0)).
\end{aligned}$$

3. $\mathbb{E}\lambda_{ij}^k = \mu_{\lambda_{ij}^k}$.

4. $\mathbb{E}z_{ij}^k = \mathbb{E}\lambda_{ij}^k + p_{ij}^k \frac{\phi(\mathbb{E}\lambda_{ij}^k)}{\Phi(\mathbb{E}\lambda_{ij}^k)} - (1 - p_{ij}^k)\frac{\phi(\mathbb{E}\lambda_{ij}^k)}{1-\Phi(\mathbb{E}\lambda_{ij}^k)}$.

   Approximation:

   When $\mathbb{E}\lambda_{ij}^k \to +\infty$, we have $\frac{\phi(\mathbb{E}\lambda_{ij}^k)}{\Phi(\mathbb{E}\lambda_{ij}^k)} \approx 0$ and $\frac{\phi(\mathbb{E}\lambda_{ij}^k)}{1-\Phi(\mathbb{E}\lambda_{ij}^k)} \approx \mathbb{E}\lambda_{ij}^k$.

   When $\mathbb{E}\lambda_{ij}^k \to -\infty$, we have $\frac{\phi(\mathbb{E}\lambda_{ij}^k)}{\Phi(\mathbb{E}\lambda_{ij}^k)} \approx -\mathbb{E}\lambda_{ij}^k$ and $\frac{\phi(\mathbb{E}\lambda_{ij}^k)}{1-\Phi(\mathbb{E}\lambda_{ij}^k)} \approx 0$.

5. $\mathbb{E}[\omega_{ii}^k] = \frac{n_k/2 + a_\omega}{\frac{1}{2}\mathbb{E}\|Y_i(\mathbb{S}^k) - \sum_{j \neq i}^p H_{ij}^k \cdot \tilde{v}_{ij}^k \cdot I(z_{ij}^k > 0)\|^2 + b_\omega}$.

6. $\mathbb{E}[R_{ij}^k] = Y_i(\mathbb{S}^k) - \sum_{j' \neq i,j} G_{ij'k}$.

#### S1.8 Algorithm

**Algorithm 1** Variational inference for `mSGR`

**Input:** Data $\{Y_i(\mathbb{S}^k)\}$, $\{H_{ij}^k\}$, hyperparameters $a_\omega, b_\omega, \sigma^2_{\Lambda i}, U, m_{\Lambda i,\text{prior}}, \sigma^2_{\text{gp}}, \hat{\eta}$, damping `step` $\in (0,1]$, tolerance `tol`, max iter $T_{\max}$

**Parameters:** $\omega_{ii}^k, \Lambda_{ij}, \tilde{v}_{ij}^k, p_{ij}^k, z_{ij}^k$

Initialize all variational parameters $\theta^{(0)}$.

**while do** $\|\theta^{(t)} - \theta^{(t-1)}\|_\infty >$ `tol` & $t \le T_{\max}$ do

  **for** $k = 1, \dots, K$ **do**

    **for** $i = 1, \dots, p$ **do**

      **1. Update** $\mathbb{E}_q(\omega_{ii}^k)$ with

      $$\omega_{ii}^k \sim \text{Gamma}\left(\frac{N_k}{2} + a_\omega,\ \frac{1}{2}\,\mathbb{E}\left\|Y_i(\mathbb{S}^k) - \textstyle\sum_{j\neq i} H_{ij}^k \cdot \tilde{v}_{ij}^k \cdot I(z_{ij}^k > 0)\right\|^2 + b_\omega\right)$$

      **for** $j \neq i$ **do**

        **2. Update** $\mathbb{E}_q(\Lambda_{ij})$ with

        $$\Lambda_{ij} \sim \mathcal{N}\left[\left(I + \frac{1}{\sigma^2_{\Lambda i}}U^{-1}\right)^{-1}\left(\mathbb{E}Z_{ij} + \frac{1}{\sigma^2_{\Lambda i}}U^{-1}m_{\Lambda i,\text{prior}}\right), \left(I + \frac{1}{\sigma^2_{\Lambda i}}U^{-1}\right)^{-1}\right]$$

        **3. Update** $\mathbb{E}_q(\tilde{v}_{ij}^k \mid z_{ij}^k)$ with $\tilde{v}_{ij}^k \mid z_{ij}^k \sim \mathcal{MVN}\left(\hat{\mu}_{\tilde{v}}(z_{ij}^k), \hat{\Sigma}_{\tilde{v}}(z_{ij}^k)\right)$

        $$\hat{\Sigma}_{\tilde{v}}(z_{ij}^k) = \left[\mathbb{E}\omega_{ii}^k I(z_{ij}^k > 0)\|H_{ij}^k\|^2 + \mathbb{E}\frac{1}{\sigma^2_{gp}}\hat{\eta}^{-1}\right]^{-1}$$

        $$\hat{\mu}_{\tilde{v}}(z_{ij}^k) = \hat{\Sigma}_{\tilde{v}}(z_{ij}^k) \cdot \mathbb{E}\omega_{ii}^k\left[I(z_{ij}^k > 0)\right](H_{ij}^k)^T \mathbb{E}R_{ij}^k$$

        **4. Update** $\mathbb{E}z_{ij}^k$ with

        $$z_{ij} \sim p_{ij}^k \cdot TN(\mathbb{E}\lambda_{ij}^k, 1, 0, +\infty) + (1 - p_{ij}^k) \cdot TN(\mathbb{E}\lambda_{ij}^k, 1, -\infty, 0)$$

        **5. Update** $p_{ij}^k$ with

        $$\text{logit}(p_{ij}^k) = \frac{1}{2}\left[\log\det(\hat{\Sigma}_{\tilde{v}}(z_{ij}^k > 0)) - \log\det(\hat{\Sigma}_{\tilde{v}}(z_{ij}^k < 0))\right] + \frac{1}{2}\hat{\mu}_{\tilde{v}}^T(z_{ij}^k > 0)\hat{\Sigma}_{\tilde{v}}^{-1}(z_{ij}^k > 0)\hat{\mu}_{\tilde{v}}(z_{ij}^k > 0) + \text{logit}(\Phi(\mathbb{E}\lambda_{ij}^k))$$

        Approximation is used when $\mathbb{E}\lambda_{ij}^k$ has large positive or negative value

      **end for**

    **end for**

  **end for**

**Damping (relaxation) update**

  Collect current parameters into $\theta^{(t)}$.

  Apply $\theta^{(t)} \leftarrow \texttt{step} \cdot \theta^{(t)} + (1 - \texttt{step}) \cdot \theta^{(t-1)}$.

**Convergence check**

  **if** $\|\theta^{(t)} - \theta^{(t-1)}\|_\infty <$ `tol` **then**

    **stop**

  **end if**

**end while**

**return** $\theta$.

#### S1.9 Postprocessing Steps

**Selection threshold determination** : As discussed in Section 3, we use a Bayesian local false discovery rate (FDR) control procedure that aims to control the average Bayesian FDR at a pre-specified level $\alpha$ (Baladandayuthapani et al., 2010) to decide selection threshold $\kappa_\alpha$. Define the Bayesian q-value (or local FDR estimate) for each edge as $\hat{q}_{ij}^k = \hat{q}_{ji}^k = 1 - \hat{p}_{ij}^k$. The threshold $\kappa_\alpha$ is then obtained via the following procedure: 1) Sort the set $\{\hat{q}_{ij}^k\}_{1 \le i < j \le p,\, 1 \le k \le K}$ in ascending order and denote the ordered sequence by $\{\hat{q}^{(t)}\}_{1 \le t \le p(p-1)K/2}$; 2) Compute the cumulative average $\bar{q}^{(t)} = \frac{1}{t} \sum_{s=1}^{t} \hat{q}^{(s)}$; 3) Find the largest index $t^*$ such that $\bar{q}^{(t^*)} \le \alpha$; 4) Set the threshold $\kappa_\alpha = 1 - \hat{q}^{(t^*)}$.

**Positive definiteness property:** From the GGM property, we have $\omega_{ij}(s^k) = -\omega_{ii}^k \cdot \gamma_{ij}(s^k)$. Based on Equations 1 and 2, for a selected edge we have

$$\sum_{j \ne i}^{p} |\omega_{ij}(s^k)| \approx \omega_{ii}^k \sum_{j \ne i}^{p} \left| \sum_{l=1}^{L} u_{ijl}^k B_l(s^k) \right|.$$

To ensure diagonal dominance, it suffices to require

$$\sum_{j \ne i}^{p} |\omega_{ij}(s^k)| \le |\omega_{ii}^k|, \qquad \forall s^k \in \mathbb{S}^k,$$

which implies

$$\sum_{j \ne i}^{p} \left| \sum_{l=1}^{L} u_{ijl}^k B_l(s^k) \right| \le 1, \qquad \forall s^k \in \mathbb{S}^k.$$

Define $B(\mathbb{S}^k) := \left[ B_1(\mathbb{S}^k) \cdots B_L(\mathbb{S}^k) \right] \in \mathbb{R}^{N_k \times L}$ and let $a_k = \|B(\mathbb{S}^k)\|_\infty$, which is an observed quantity depending on the spatial domain and GP parameters. Then,

$$\sum_{j \ne i}^{p} \left| \sum_{l=1}^{L} u_{ijl}^k B_l(s^k) \right| \le a_k \sum_{j \ne i}^{p} \left| \sum_{l=1}^{L} u_{ijl}^k \right| \le a_k \sum_{j \ne i}^{p} \sum_{l=1}^{L} |u_{ijl}^k|.$$

Therefore, diagonal dominance is guaranteed if

$$a_k \sum_{j \ne i}^{p} \sum_{l=1}^{L} |u_{ijl}^k| \le 1.$$

To enforce this constraint, we rescale $u_{ijl}^k$ by the factor $a_k \sum_{j \ne i}^{p} \sum_{l=1}^{L} |u_{ijl}^k|$.

### S2 Simulation Studies

#### S2.1 Metrics for comparison

We use the following measures to compare with other methods (I) true positive rate (TPR); (II) false positive rate (FPR); (III) false discovery rate (FDR); (IV) Matthews correlation coefficient (MCC). MCC (Matthews, 1975) measures the quality of binary classification, ranging from +1 (perfect classification) to -1 (total mismatch).

#### S2.2 Comparative Methods

We compare `mSGR` with the following methods (I) Spatial Graphical Regression (SGR, Acharyya et al. (2025)); (II) GraphR (Chen et al., 2025); (III) Joint Graphical Lasso (JGL, Danaher et al. (2014)); (IV) jointGHS (Lingjærde et al., 2024) and (V) graphical lasso (Glasso, Friedman et al. (2008)) method applied independently to each FOV.

- **SGR**: The original `SGR` method is implemented using a Markov chain Monte Carlo (MCMC) algorithm. To ensure a fair computational comparison with variational methods, we derive a MFVB approximation for `SGR` and implement it using the `mSGR` package. Hyperparameters are set to $a_\omega = b_\omega = 10$, and the prior edge inclusion probability $\pi$ is fixed to match the true sparsity level in each simulation setting. For GP parameters, we set $\sigma^2_{gp} = 1, a_{gp} = 0.01, b_{gp} = 0.5$, approximation degree $= 10$. The MFVB algorithm is run until convergence using the same learning rate and stopping criteria as `mSGR`.

- **GraphR**: The algorithm is implemented using the `GraphR` package with hyperparameters set to $a_\pi = 1$, $b_\pi = 4$, and $a_\tau = b_\tau = 0.005$, with a maximum of 2000 iterations and a convergence tolerance of $10^{-3}$. We apply `GraphR` separately to each FOV, using scaled $(x, y)$ coordinates as continuous external covariates. Since `GraphR` outputs cell-level edge inclusion probabilities (rather than FOV-level summaries), we construct an FOV-level edge indicator by declaring an edge present in a given FOV if it is selected in at least one cell within that FOV.

- **GGL**: The tuning parameters $\lambda_1$ and $\lambda_2$ are selected from a $20 \times 20$ grid, evenly spaced between 0.05 and 0.75 for both $\lambda_1$ and $\lambda_2$, using the approximated Akaike Information Criterion (AIC). Each FOV is treated as a distinct group, and spatial coordinate information is not incorporated in this method.

- **jointGHS**: The algorithm is implemented using the `jointGHS` package with `epsilon` $= 0.001$ and `AIC_eps` $= 0.001$. Similar to GGL, each FOV is regarded as a separate group, and no spatial coordinate information is used in model fitting.

- **GLASSO**: The tuning parameter $\lambda$ is selected using the stability-based approach Liu et al. (2010). The graphical lasso model is fit separately for each FOV, and no spatial coordinate information is incorporated.

#### S2.3 Additional Simulation Results of Scenario I

We compare mSGR with competing methods under alternative sparsity levels (10%). All methods are implemented using the same parameter settings as in Section 4 to ensure a fair comparison. mSGR consistently outperforms or matches competing approaches in terms of graph recovery, demonstrating its robustness to increased network density.

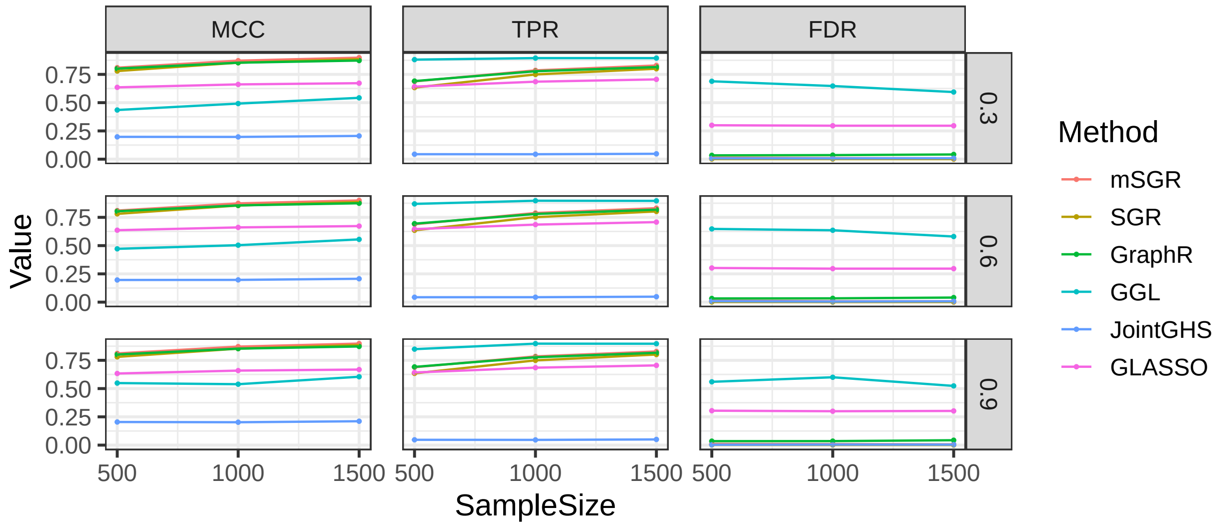


Figure S1: **Simulation performance of mSGR.** Simulation results comparing different methods under varying correlation strengths between FOVs ($\rho$ = 0.3, 0.6, 0.9) and sample sizes ($n = 500, 1000, 1500$ per FOV) with sparsity fixed to be 10%. The plot represents mean values of the evaluation metrics (MCC, TPR, FDR, FPR) across replications.

We evaluate the robustness of mSGR to the choice of Gaussian process hyperparameters by varying $a_{gp}$ and $b_{gp}$ over a wide range of values under different sparsity setting (5% and 10%) as shown in Figures S1 − S7. Across all configurations, mSGR exhibits stable performance in terms of edge recovery, with minimal sensitivity to the specific hyperparameter settings. These results suggest that the proposed method is robust to moderate misspecification of the GP prior.

#### S2.4 Computation Time

Figure S8 summarizes the average computation time of all competing methods in Scenario I. Computation time increases with sample size for all methods, while remaining relatively in-

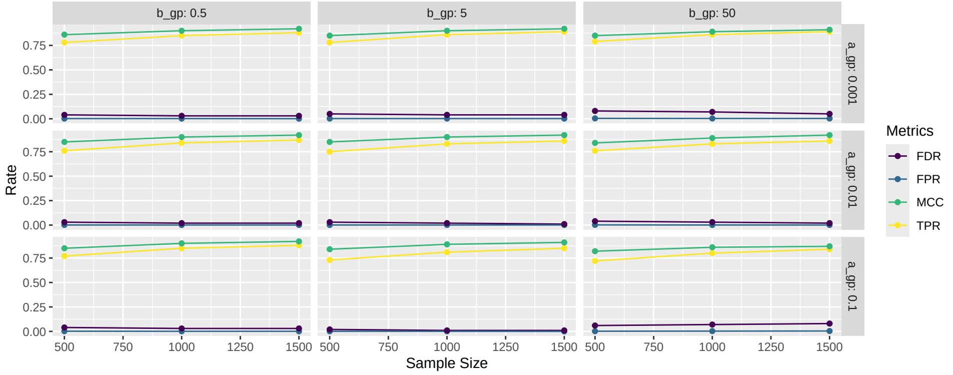


Figure S2: Simulation results comparing different hyperparameter settings, with $a_{gp} \in 0.001, 0.01, 0.1$ and $b_{gp} \in 0.5, 5, 50$. The correlation strength between FOVs is fixed at 0.3, sparsity is set to 5%, and the per-FOV sample size varies across $n \in 500, 1000, 1500$.

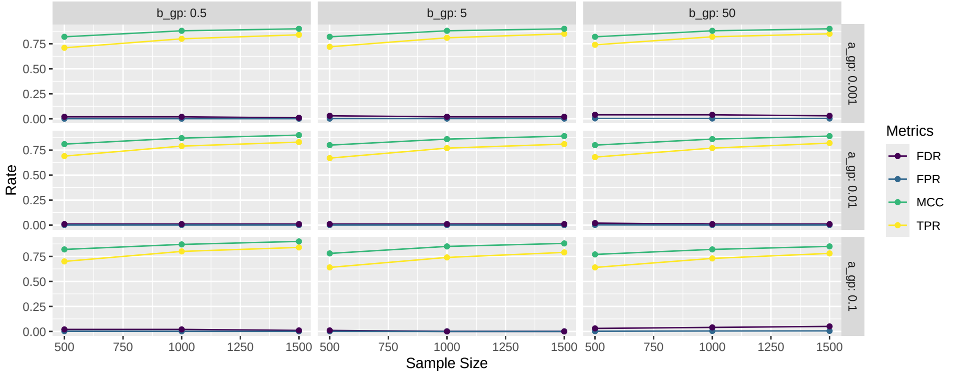


Figure S3: Simulation results comparing different hyperparameter settings, with $a_{gp} \in 0.001, 0.01, 0.1$ and $b_{gp} \in 0.5, 5, 50$. The correlation strength between FOVs is fixed at 0.3, sparsity is set to 10%, and the per-FOV sample size varies across $n \in 500, 1000, 1500$.

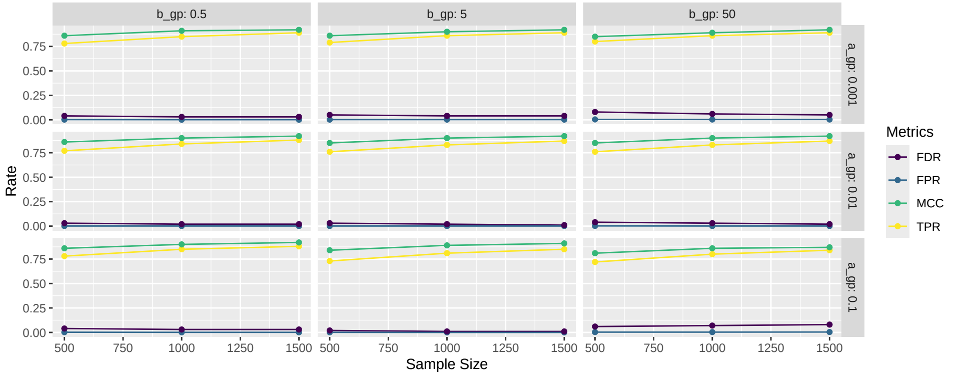


Figure S4: Simulation results comparing different hyperparameter settings, with $a_{gp} \in 0.001, 0.01, 0.1$ and $b_{gp} \in 0.5, 5, 50$. The correlation strength between FOVs is fixed at 0.6, sparsity is set to 5%, and the per-FOV sample size varies across $n \in 500, 1000, 1500$.

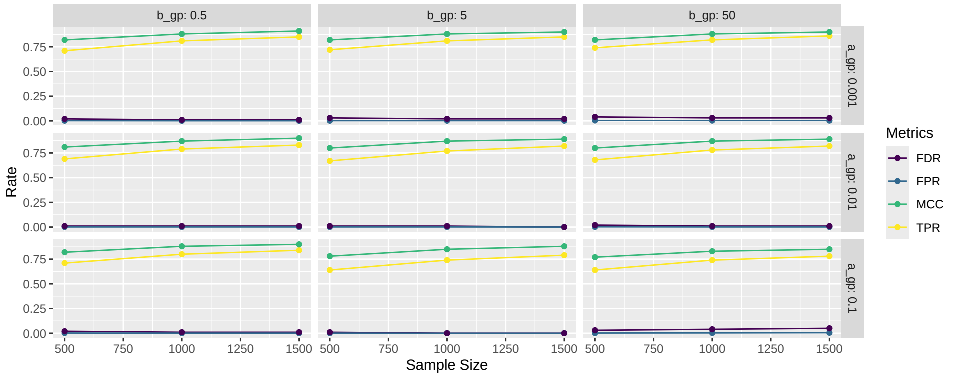


Figure S5: Simulation results comparing different hyperparameter settings, with $a_{gp} \in 0.001, 0.01, 0.1$ and $b_{gp} \in 0.5, 5, 50$. The correlation strength between FOVs is fixed at 0.6, sparsity is set to 10%, and the per-FOV sample size varies across $n \in 500, 1000, 1500$.

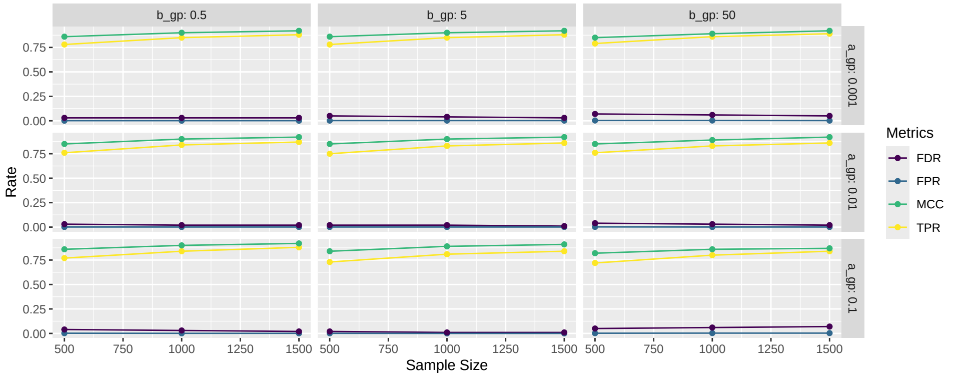


Figure S6: Simulation results comparing different hyperparameter settings, with $a_{gp} \in 0.001, 0.01, 0.1$ and $b_{gp} \in 0.5, 5, 50$. The correlation strength between FOVs is fixed at 0.9, sparsity is set to 5%, and the per-FOV sample size varies across $n \in 500, 1000, 1500$.

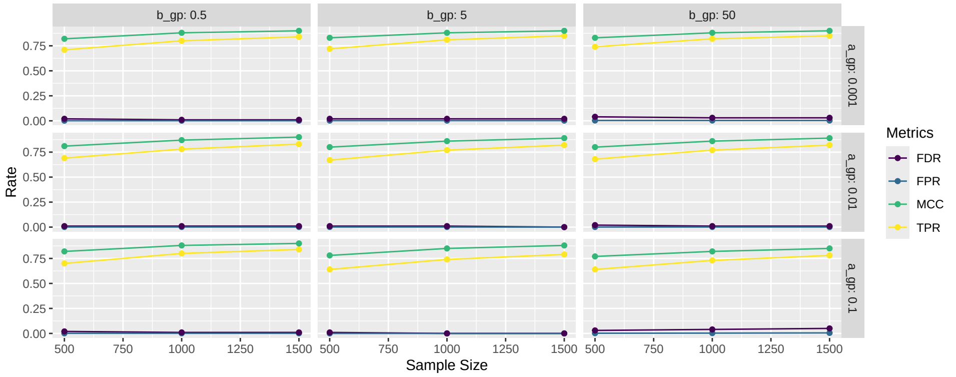


Figure S7: Simulation results comparing different hyperparameter settings, with $a_{gp} \in 0.001, 0.01, 0.1$ and $b_{gp} \in 0.5, 5, 50$. The correlation strength between FOVs is fixed at 0.9, sparsity is set to 10%, and the per-FOV sample size varies across $n \in 500, 1000, 1500$.

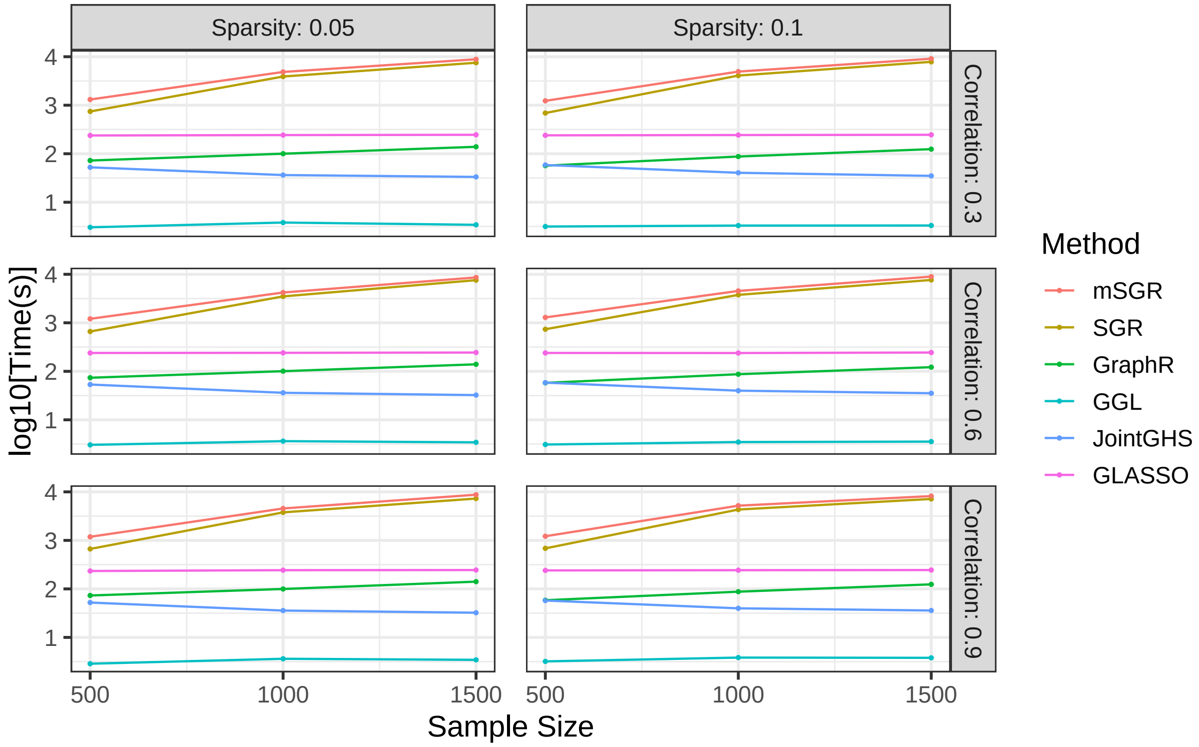


Figure S8: **Computation time comparison across methods for Scenario I.** Average computation time (log10 scale) as a function of sample size per FOV for `mSGR`, SGR, GraphR, GGL, JointGHS, and GLASSO. Results are shown for sparsity levels 0.05 and 0.1 (columns) and inter-FOV correlation strengths $\rho = 0.3, 0.6, 0.9$ (rows). Computation time is averaged across replicates and reported in second before log transformation.

sensitive to sparsity level and inter-FOV correlation strength. For `mSGR`, the average runtime per replicate increases from approximately 0.34 hours at $n = 500$ to 1.26 hours at $n = 1000$ and 2.43 hours at $n = 1500$ per FOV, with similar trends observed across correlation and sparsity settings. Table S1 reports the average runtime per simulation replicate for each method in Scenario II, measured in minutes under the same computational environment using a single CPU core on a Linux-based HPC cluster.

### S3 Spatial network characterization of EMT gradient in sarcomatoid renal cancer

#### S3.1 Expression normalization

To prepare the gene expression data for spatial analysis, we exclude cells with total counts less than 10, resulting in 28,257 remaining cells. The raw count matrix is then normalized

| Method | Time (min) |
|---|---|
| **mSGR** | 80.17 |
| SGR | 57.19 |
| GraphR | 0.95 |
| GGL | 0.05 |
| JointGHS | 0.47 |
| GLASSO | 3.05 |

Table S1: Average computation time per simulation replicate (in minutes) in Scenario II.

using the SCTransform procedure implemented in the Seurat framework (Hafemeister and Satija, 2019). After applying normalization, the distribution of normalized gene expression counts exhibited an approximately Gaussian-like shape.

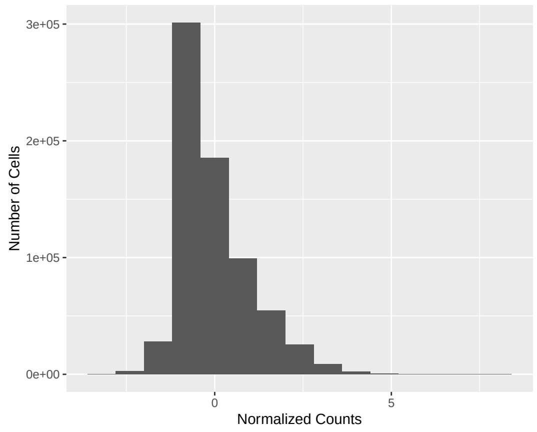


Figure S9: Distribution of normalized gene expression

#### S3.2 Incorporation of spatial coordinates at FOV-level:

We consider a power correlation matrix with $[k, k']$ element defined as $\rho^{d_{kk'}}$, where $d_{kk'}$ denotes the Euclidean distance between the centroids of FOVs $k$ and $k'$. To estimate $\rho$, we first aggregate cell-level expression data to the FOV level. Specifically, for each FOV we compute the mean expression of all EMT genes, which serves as the variable of interest for calculating spatial autocorrelation. Pairwise Euclidean distances between FOV centroids are then used to construct a Gaussian kernel spatial weight matrix with entries

$$w_{kk'} = \exp\left(-\frac{d_{kk'}^2}{2\sigma^2}\right),$$

where $\sigma = 1$ serves as the bandwidth parameter. The spatial autocorrelation parameter $\rho$ is quantified using Moran's $I$ statistic, with the Gaussian kernel matrix serving as the spatial weight structure.

**Incorporation of spatial coordinates at cell-level:** Prior to fitting the Gaussian process, the $x$ and $y$ coordinates of all cells are scaled to ensure comparable ranges and numerical stability.

#### S3.3 Hyperparameters Specification

For the selection indicators, we specify the prior hyperparameters as $M = \mathbf{0}, U = I_{23}, V = \left[V_{kk'}\right]_{23\times 23}, V_{kk'} = \frac{1}{50}\rho^{d_{kk'}}$. Here, $d_{kk'}$ denotes the Euclidean distance between the centroids of FOVs $k$ and $k'$, and $\rho^{d_{kk'}}$ defines the power correlation kernel applied elementwise. The Gaussian process prior is controlled by shape parameters $a_{\text{gp}} = 0.01$ and $b_{\text{gp}} = 0.5$, with polynomial degree set to be 10. We set $a_w = 10$ and $b_w = 10$. Convergence is declared once the magnitude of the difference between updated and previous parameter estimates falls below 0.001.

#### S3.4 Connectivity Between Genes in EMT Pathways

We provide additional results characterizing both connectivity degree and gene expression patterns for genes that exhibit statistically significant differences in connectivity degree along the tumor progression gradient or are identified as hub genes. Specifically, we present gene-level connectivity summaries together with corresponding expression levels, highlighting genes whose network involvement varies markedly across spatial contexts.

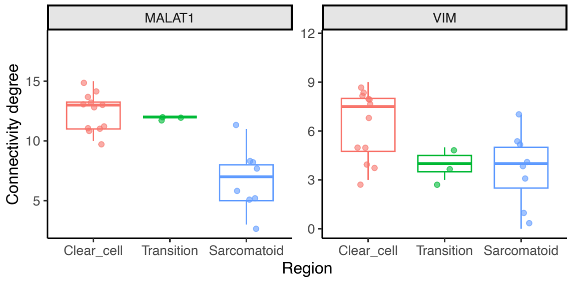


Figure S10: **MALAT1 and VIM as hub genes with high connectivity scores.** Boxplots show the distribution of FOV connectivity degree for MALAT1 and VIM across clear cell, transitional, and sarcomatoid regions. Connectivity degree is defined as the number of significant gene–gene associations incident to each gene within a given FOV. Each point represents one FOV.

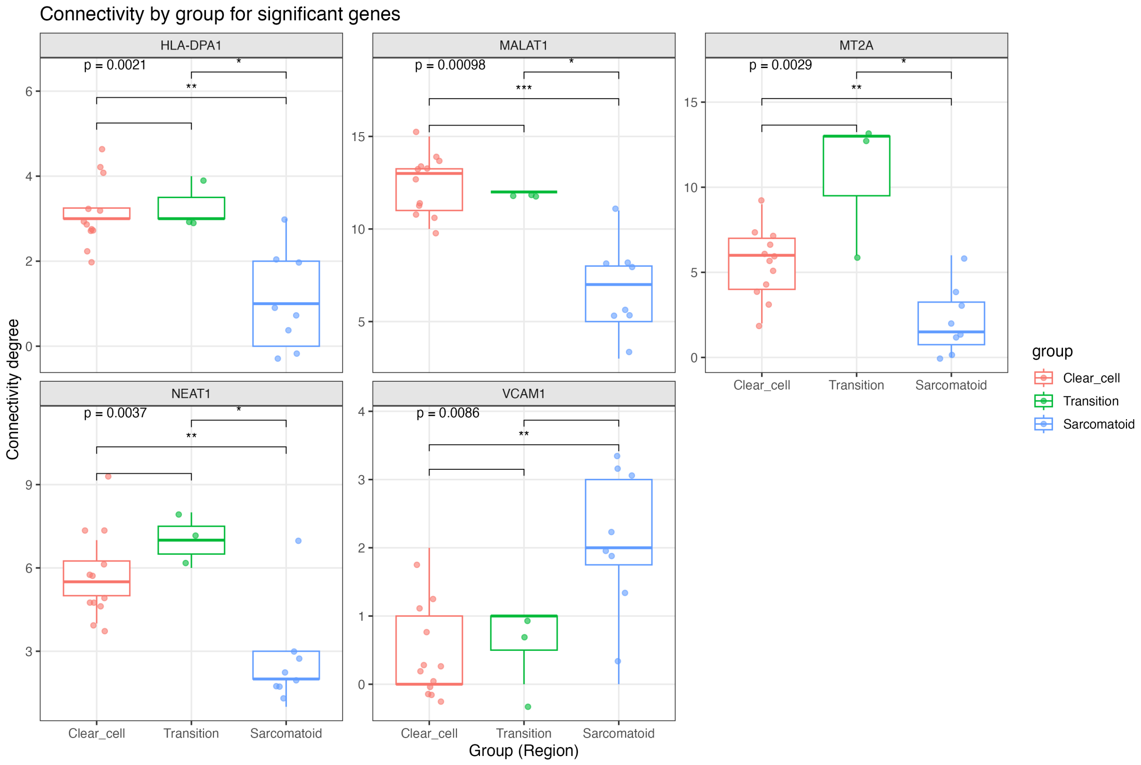


Figure S11: **Regional differences in gene connectivity for HLA-DPA1, MALAT1, MT2A, NEAT1 and VCAM1** Boxplots show the distribution of FOV connectivity degree for five genes (HLA-DPA1, MALAT1, MT2A, NEAT1 and VCAM1) which show significant regional differences after multiple testing correction. Connectivity degree is defined as the number of significant gene–gene associations incident to each gene within a given FOV. Each point represents one FOV. Overall differences across regions were assessed using the Kruskal–Wallis test, followed by pairwise Wilcoxon rank-sum tests with significance indicated by stars (* $p < 0.05$, ** $p < 0.01$, *** $p < 0.001$)

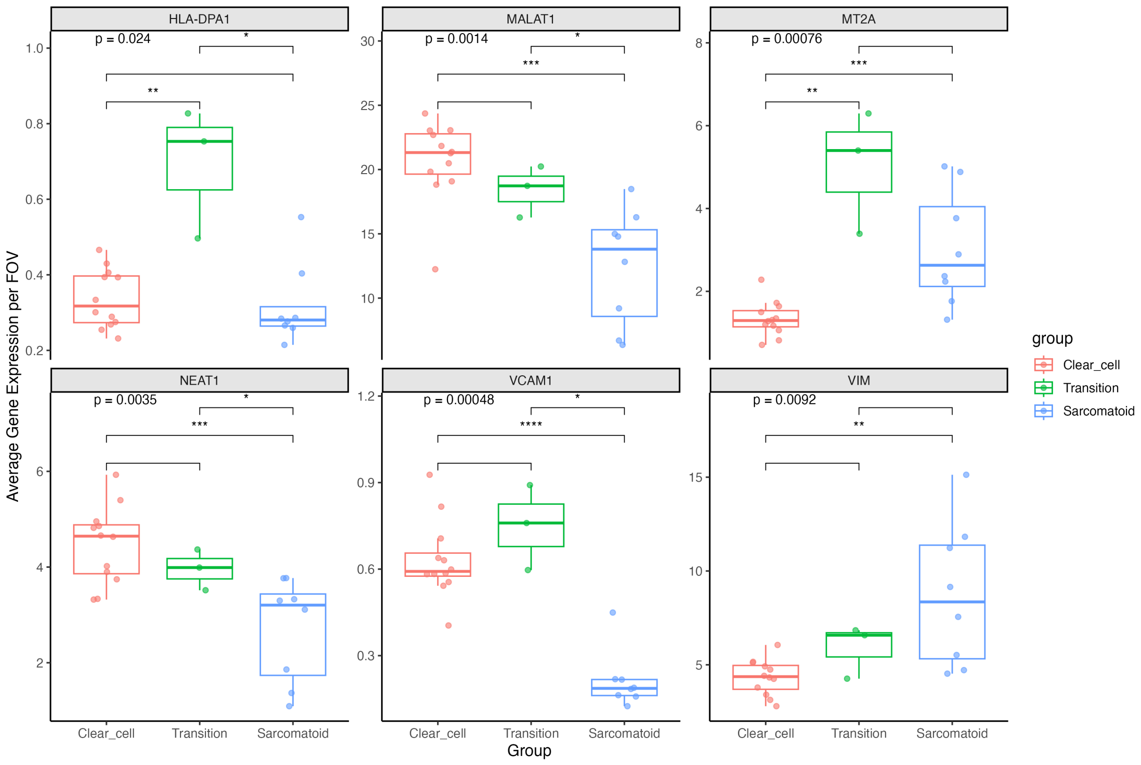


Figure S12: **Regional differences in gene expression for HLA-DPA1, MALAT1, MT2A, NEAT1, VCAM1 and VIM.** Boxplots display the distribution of mean gene expression at the FOV level for six genes (HLA-DPA1, MALAT1, MT2A, NEAT1, VCAM1 and VIM). Each point corresponds to a single FOV. Overall differences across regions were evaluated using the Kruskal–Wallis test, followed by pairwise Wilcoxon rank-sum tests, with statistical significance indicated by stars (* $p < 0.05$, ** $p < 0.01$, *** $p < 0.001$).

### References


Acharyya, S., Kang, J., and Baladandayuthapani, V. (2025). Spatially varying graph estimation for spatial transcriptomics cancer data. *bioRxiv*, pages 2025–05.

Baladandayuthapani, V., Ji, Y., Talluri, R., Nieto-Barajas, L. E., and Morris, J. S. (2010). Bayesian random segmentation models to identify shared copy number aberrations for array cgh data. *Journal of the american statistical association*, 105(492):1358–1375.

Banerjee, A., Dunson, D. B., and Tokdar, S. T. (2013). Efficient gaussian process regression for large datasets. *Biometrika*, 100(1):75–89.

Banerjee, S., Carlin, B. P., and Gelfand, A. E. (2003). *Hierarchical modeling and analysis for spatial data*. Chapman and Hall/CRC.

Chen, L., Acharyya, S., Luo, C., Ni, Y., and Baladandayuthapani, V. (2025). A probabilistic modeling framework for genomic networks incorporating sample heterogeneity. *Cell Reports Methods*, 5(2).

Cressie, N. and Wikle, C. K. (2011). *Statistics for spatio-temporal data*. John Wiley & Sons.

Danaher, P., Wang, P., and Witten, D. M. (2014). The joint graphical lasso for inverse covariance estimation across multiple classes. *Journal of the Royal Statistical Society: Series B (Statistical Methodology)*, 76(2):373–397.

Friedman, J., Hastie, T., and Tibshirani, R. (2008). Sparse inverse covariance estimation with the graphical lasso. *Biostatistics*, 9(3):432–441.

Hafemeister, C. and Satija, R. (2019). Normalization and variance stabilization of single-cell rna-seq data using regularized negative binomial regression. *Genome biology*, 20(1):296.

Lin, Z., Si, Y., and Kang, J. (2024). Latent subgroup identification in image-on-scalar regression. *The annals of applied statistics*, 18(1):468.

Lingjærde, C., Fairfax, B. P., Richardson, S., and Ruffieux, H. (2024). Scalable multiple network inference with the joint graphical horseshoe. *The Annals of Applied Statistics*, 18(3):1899 – 1923.

Liu, H., Roeder, K., and Wasserman, L. (2010). Stability approach to regularization selection (stars) for high dimensional graphical models. *Advances in neural information processing systems*, 23.

Matthews, B. W. (1975). Comparison of the predicted and observed secondary structure of t4 phage lysozyme. *Biochimica et Biophysica Acta-Protein Structure*, 405(2):442–451.

Minh, H. Q., Niyogi, P., and Yao, Y. (2006). Mercer's theorem, feature maps, and smoothing. In *International Conference on Computational Learning Theory*, pages 154–168. Springer.

Shi, R. and Kang, J. (2015). Thresholded multiscale gaussian processes with application to bayesian feature selection for massive neuroimaging data. *arXiv preprint arXiv:1504.06074*.

Wang, L. (2008). *Karhunen-Loeve expansions and their applications.* London School of Economics and Political Science (United Kingdom).
